# Selective regulation of IFN-γ and IL-4 co-producing unconventional T cells by purinergic signalling

**DOI:** 10.1101/2024.08.11.607476

**Authors:** Calvin Xu, Andreas Obers, Minyi Qin, Alice Brandli, Joelyn Wong, Xin Huang, Allison Clatch, Aly Fayed, Graham Starkey, Rohit D’Costa, Claire L Gordon, Lynette Beattie, Laura K. Mackay, Dale I. Godfrey, Hui-Fern Koay

**Affiliations:** Department of Microbiology and Immunology, The University of Melbourne at The Peter Doherty Institute for Infection and Immunity, Melbourne, Victoria, Australia; Department of Anatomy and Physiology, The University of Melbourne, Melbourne, Victoria, Australia; The Florey Institute of Neuroscience and Mental Health, Melbourne, Victoria, Australia; The State Key Laboratory of Pharmaceutical Biotechnology, School of Life Sciences, Nanjing University, Nanjing, Jiangsu; Liver & Intestinal Transplant Unit, Austin Health, Melbourne, VIC, Australia; Department of Surgery, The University of Melbourne, Austin Health, Melbourne, VIC, Australia; DonateLife Victoria, Carlton, Victoria; Department of Intensive Care Medicine, Melbourne Health, Melbourne, VIC, Australia; Department of Infectious Diseases, Austin Health, Melbourne, VIC, Australia; North Eastern Public Health Unit, Austin Health, Melbourne, VIC, Australia

## Abstract

Unconventional T cells, including mucosal-associated invariant T (MAIT), natural killer T (NKT), and gamma-delta T (γδT) cells, comprise distinct T-bet^+^, IFN-γ^+^ and RORγt^+^, IL-17^+^ subsets which play differential roles in health and disease. NKT1 cells are susceptible to ARTC2-mediated P2X7 receptor (P2RX7) activation, but the effects on other unconventional T-cell types are unknown. Here, we show that MAIT, γδT, and NKT cells express P2RX7 and are sensitive to P2RX7-mediated cell death. Mouse peripheral T-bet^+^ MAIT1, γδT1, and NKT1 cells, especially in liver, co-express ARTC2 and P2RX7, which can be further upregulated by retinoic acid. Blocking ARTC2 or inhibiting P2RX7 protected MAIT1, γδT1, and NKT1 cells from cell death, enhanced their survival *in vivo*, and increased the number of IFN-γ-secreting cells without affecting IL-17 production. Importantly, this revealed the existence of IFN-γ and IL-4 co-producing unconventional T-cell populations normally lost upon isolation due to ARTC2/P2RX7-induced death. Administering extracellular NAD *in vivo* activated this pathway, depleting P2RX7-sensitive unconventional T cells. Our study reveals ARTC2/P2RX7 as a common regulatory axis modulating the unconventional T-cell compartment, affecting the viability of IFN-γ- and IL-4-producing T cells, offering important insights to facilitate future studies into how these cells can be regulated in health and disease.

## Introduction

Unconventional T-cell lineages, including MR1-restricted mucosal-associated invariant T (MAIT) cells, CD1d-restricted natural killer T (NKT) cells, and gamma-delta T (γδT) cells, are characterised by their specificity for non-peptide antigens, and rapid and potent cytokine production immediately upon activation (Godfrey et al., 2015). Their ‘innate-like’ effector function is imbued during thymic development and is thought to be dependent on the master transcription factor, promyelocytic leukemia zinc finger (PLZF) (Koay et al., 2016; Kreslavsky et al., 2009; Lu et al., 2015; Pellicci et al., 2020; Savage et al., 2008). Developing MAIT, NKT, and some γδT cells differentially switch on other lineage-defining transcription factors, including T-bet, RORγt, and GATA-3, that drive their differentiation into mature and functionally distinct subsets (Mayassi et al., 2021; Pellicci et al., 2020). Generally, T-bet^+^ MAIT1, γδT1, and NKT1 cells produce IFN-γ, whilst RORγt^+^ MAIT17, γδT17, and NKT17 cells produce IL-17 (Lee et al., 2020). Further highlighting their functional differences, MAIT1, γδT1, and NKT1 cells preferentially accumulate in tissues such as liver and spleen, while their RORγt^+^ counterparts are enriched within lymph nodes, lungs, and skin (Chandra et al., 2023; Crosby and Kronenberg, 2018; Ribot et al., 2021; Salou et al., 2019). Though a subset of γδT and NKT cells can express GATA-3 and mainly produce IL-4, many γδT and NKT cells express GATA-3 in addition to T-bet, and characteristically co-produce IFN-γ and IL-4 (Azuara et al., 1997; Gerber et al., 1999; Lee et al., 2013; Lee et al., 2015; Narayan et al., 2012; Pereira et al., 2013). Similarly, RORγt^+^ NKT17 cells can also express GATA-3 and co-produce IL-17 and IL-4 (Cameron and Godfrey, 2018). Whilst MAIT cells are known to potently produce IL-17 and IFN-γ upon activation, a clear subset of IL-4-producing MAIT cells has remained elusive (Lee et al., 2020; Pellicci et al., 2020; Salou et al., 2019) despite low levels of IL-4 detected in the supernatants of stimulated mouse MAIT cells (Rahimpour et al., 2015) and chronically stimulated human MAIT cells (Kelly et al., 2019). As functionally distinct subsets of MAIT, γδT, and NKT cells play different and often opposing roles in the immune response (Mayassi et al., 2021), understanding the mechanisms that govern their diverse activity will provide insight into improving future immunotherapies that target these cells.

MAIT, γδT, and NKT cells are particularly abundant in liver, in which they comprise up to 50% of T cells and are biased towards T-bet^+^, IFN-γ-producing functional subsets (Matsuda et al., 2000; Salou et al., 2019; Xu et al., 2023). As the liver plays a central role in both immune defence and nutrient metabolism, amongst other roles, where gut-derived dietary and microbial antigens are brought to the liver via the portal vein, immune homeostasis in this organ must balance tolerance to innocuous antigens with resistance to pathogens (Protzer et al., 2012). Purinergic signaling may represent a regulatory mechanism here, where the detection of purine nucleotides and related-compounds, including ATP and nicotinamide adenine dinucleotide (NAD), is mediated by purinergic receptors widely expressed on immune cells (Rissiek et al., 2013; Rissiek et al., 2014; Rivas-Yáñez et al., 2020). The release of intracellular ATP and NAD by stressed and/or damaged cells into the extracellular space can be detected by NKT cells via the purinergic P2X7 receptor (P2RX7) (Bovens et al., 2020; Rissiek et al., 2013; Rissiek et al., 2014). Whilst strongly activated hepatic NKT cells can mediate liver injury, these cells are, in turn, regulated by tissue damage following liberation of NAD and ATP and consequent P2RX7 activation (Bovens et al., 2020; Kawamura et al., 2006).

In contrast to direct activation by ATP, P2RX7 can be indirectly activated by NAD in a manner dependent on the ecto-ADP-ribosyltransferase, ARTC2.2 (ARTC2) (Rissiek et al., 2015; Rivas-Yáñez et al., 2020). Extracellular NAD is a substrate for ARTC2, providing an ADP-ribosyl group for ARTC2-mediated ADP-ribosylation of P2RX7, resulting in P2RX7 activation (Seman et al., 2003). Although ARTC2-mediated P2RX7 activation is well characterised in mouse models, ARTC2 is not expressed in humans (Haag et al., 1994). Other ADP-ribosyltransferases, such as ARTC1 and ARTC5, have ADP-ribosyltransferase activity, though it is unclear whether they contribute to the NAD-mediated activation of human P2RX7 (Laing et al., 2011; Leutert et al., 2018) (Cortés-Garcia et al., 2016; Hesse et al., 2022; Wennerberg et al., 2022). ATP- and NAD-induced P2RX7 activation both result in similar cellular and biochemical outcomes, where only brief exposures to low concentrations of NAD is needed to robustly drive P2RX7 activation and cell death (Rissiek et al., 2014). Accordingly, this is termed NAD-induced cell death (NICD) (Seman et al., 2003), where P2RX7 activation is also associated with the loss of the co-stimulatory molecule, CD27, from the cell surface, and reduced production of IFN-γ and IL-4 by NKT cells after stimulation (Borges Da Silva et al., 2019; Kawamura et al., 2006; Liu and Kim, 2019; Rissiek et al., 2013). Though ARTC2 and P2RX7 are differentially expressed on NKT-cell functional subsets (Borges Da Silva et al., 2019; Liu and Kim, 2019) it is unclear whether ARTC2-mediated P2RX7 activation affects specific cytokine-producing NKT-cell subsets. Furthermore, it is unknown whether this axis regulates MAIT and γδT cells, and the homeostasis of their functional subsets.

Here, we show that human liver and blood MAIT, γδT, and NKT cells express P2RX7 at higher levels relative to conventional T cells. Within mouse models, peripheral T-bet^+^ MAIT1, γδT1, and NKT1 cells, and other PLZF^+^ αβT-cell subsets co-expressed high levels of ARTC2 and P2RX7 in liver and, to a lesser extent, spleen. ARTC2 and P2RX7 expression could be further upregulated in the presence of retinoic acid. In response to cell damage during organ processing, multiple unconventional T-cell subsets upon cell culture exhibited a loss of surface CD27 expression and underwent NICD, in a manner dependent on P2RX7 activation and ARTC2 activity. Nanobody-mediated ARTC2 blockade or P2RX7 inhibition significantly increased the number of peripheral IFN-γ-producing, but not IL-17-producing, unconventional T cells upon stimulation *ex vivo*. Importantly, this increase was predominantly attributable to increased IL-4 and IFN-γ co-producing unconventional and other T-cell types, especially within liver, suggesting that this functionally distinct subset may be restrained by NAD released by damaged cells. Lastly, we show that intravenous administration of exogenous NAD rapidly depleted liver PLZF^+^T-bet^+^ unconventional T cells *in vivo*, including poorly characterised PLZF^+^ CD4^+^ and PLZF^+^ CD4^-^CD8^-^ DN non-MAIT/NKT αβT-cell subsets, in an ARTC2-dependent manner. Overall, our findings highlight the regulation of T-bet^+^ unconventional T cells and other PLZF^+^T-bet^+^ αβT-cell subsets by NAD-driven, ARTC2-mediated P2RX7 activation, where this axis may collectively modulate these cells in the context of tissue damage and disease.

## Results

### Human unconventional T cells express P2RX7

Analysis of a transcriptomic dataset (Gutierrez-Arcelus et al., 2019) on the four major unconventional T-cell populations – MAIT, NKT, V81^+^ γδT, and V82^+^ γδT cells, indicated that all these cell types express the *P2RX7* gene (**Fig. S1A**). Indeed, human γδT and MAIT cells were reported to express P2RX7 and γδT cells were susceptible to ATP-mediated P2RX7 activation and cell death (Winzer et al., 2022). We examined P2RX7 protein expression on MR1-5-OP-RU tetramer^+^ MAIT cells, CD1d-α-GalCer tetramer^+^ NKT cells, and γδT-cell subsets from human peripheral blood (**Fig. 1A** and **Fig. S1B**). This again demonstrated that P2RX7 is expressed by human unconventional T cells, although there was a wide range of expression levels on these cell types (**Fig. 1A** and **Fig. S1B**). P2RX7 expression was detected on both fresh blood samples and cryopreserved samples (**Fig. S1B**). Furthermore, one donor (Donor 03) appeared to lack P2RX7 expression on both monocytes and T cells (**Fig. S1B-C**). This variability may be attributed to single nucleotide polymorphisms within the highly polymorphic *P2RX7* gene locus, which are common in humans, impacting on P2RX7 protein expression and function (Fuller et al., 2009; Schäfer et al., 2022). Taken together, these data support the concept that human unconventional T cells express P2RX7. While we found an increase in P2RX7 expression amongst unconventional T cells compared to conventional T cells (**Fig. 1A** and **S1B, D**), in some individuals P2RX7 was expressed at similar levels across all T cells (**Fig. 1A** and **S1B, D**).

**Figure 1.**
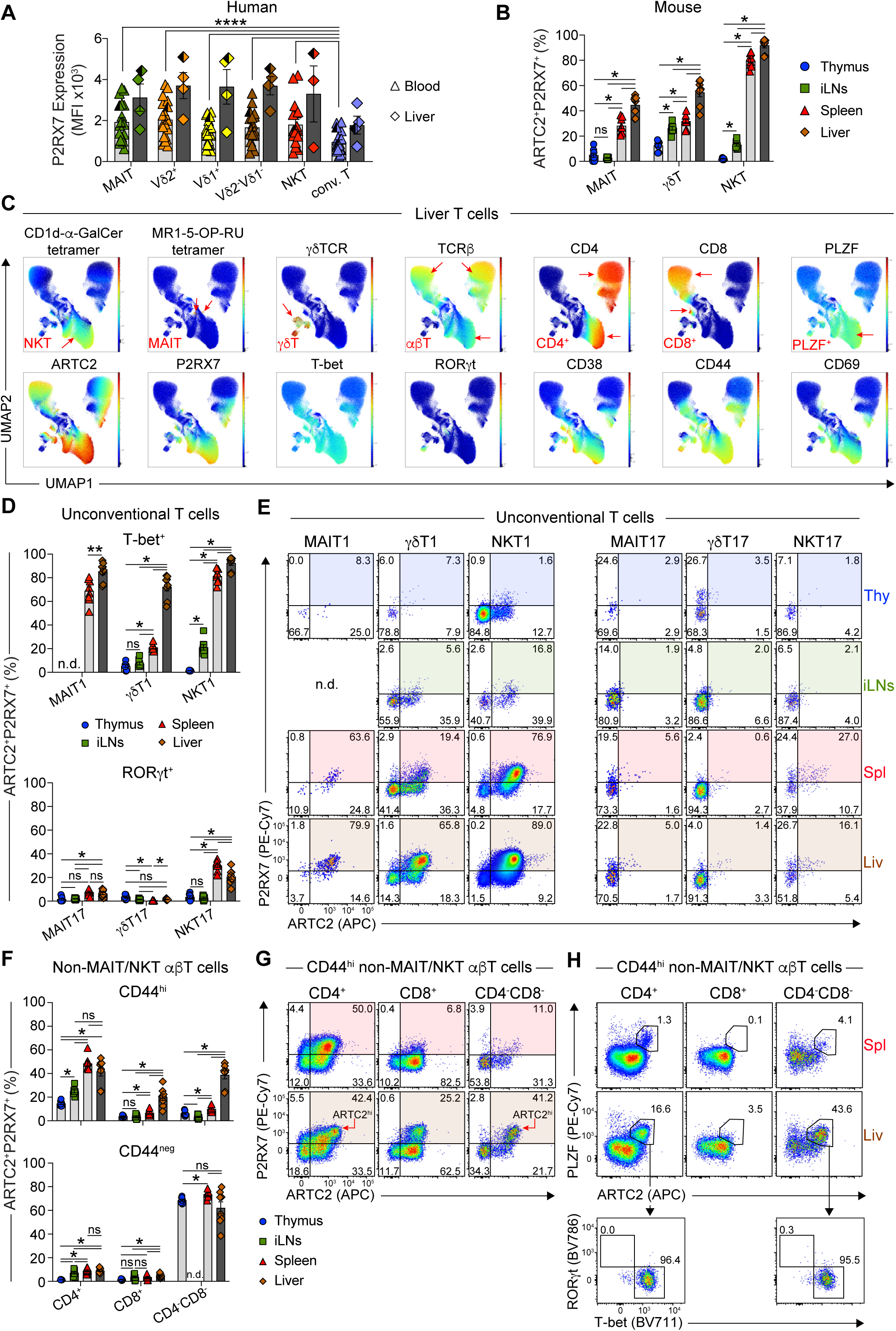
Unconventional T cells express P2RX7 to greater extent than conventional T cells. (A) Graph shows mean fluorescence intensity (MFI) of P2RX7 labelling on MAIT cells (MR1-5-OP-RU tetramer^+^CD3^+^), V82^+^, V81^+^, and V81^-^V82^-^ γδT cells (γδTCR^+^CD3^+^), NKT cells (CD1d-α-GalCer tetramer^+^CD3^+^), and conventional T cells (conv. T; defined as non-MAIT/NKT γδTCR^-^ CD3^+^ T cells) from human blood and liver. A total of 4 human liver and 18-19 blood donors were analysed across 6 separate experiments. Half-shaded symbols represent matched blood and liver samples from one donor. Due to their paucity, NKT cells within one liver and one blood donor were not analysed. (B) Graph shows percentages of ARTC2^+^P2RX7^+^ cells out of total MAIT, γδT, and NKT cells from C57BL/6 WT mouse organs. Each symbol represents an individual mouse. A total of 8 mice were analysed across 3 separate experiments. (C) UMAP representation of flow cytometric analysis of liver T cells. UMAP plots were generated by concatenation of all (n = 2) mice from one of two similar experiments. Red arrows within UMAP plots indicate various T-cell populations. (D & F) Graphs show percentages of ARTC2^+^P2RX7^+^ cells of T-bet^+^ MAIT1, γδT1, and NKT1 cells, RORγt^+^ MAIT17, γδT17, and NKT17 cells (D), and CD44^hi^ and CD44^neg^ non-MAIT/NKT αβT-cell CD4/CD8 subsets (F) from C57BL/6 WT mouse organs. (E & G) Flow cytometric analysis of ARTC2 and P2RX7 expression by indicated cell types. Red arrows in (G) indicate ARTC2^hi^ cells. (H) Flow cytometric analysis of PLZF and ARTC2, and T-bet and RORγt expression on indicated non-MAIT/NKT αβT-cell subsets. (A, B, D, F) Graphs depict individual data points and mean ± SEM. ns P>0.05, *P≤0.05, **P≤0.01, ***P≤0.001, ****P≤0.0001 using a Wilcoxon matched-pairs signed-rank test with a Bonferroni-Dunn correction for multiple comparisons where required. (E, G, H) Numbers in plots represent percentage of gated cells. (D, E, F) The percentages of ARTC2^+^P2RX7^+^ thymic MAIT1 (D), iLN MAIT1 cells (E) and iLN CD44^neg^ CD4^-^CD8^-^ T cells (F) could not be reliably determined (n.d.) due to their paucity. (E) FACS plots generated by concatenation of all (n = 3) mice from one of three similar experiments. With the exception of (H), all mice were injected with the anti-ARTC2 nanobody (clone: s+16) prior to organ harvest.

We next examined human liver samples from the Australian Donation and Transplantation Biobank. Similar to blood, considerable inter-donor variability of P2RX7 expression was observed on all liver T-cell types, with liver unconventional T cells, on average, labelling at a higher intensity than conventional T cells (**Fig. 1A**), including CD4^+^ and CD8^+^ conventional T-cell subsets defined by CD45RA, CD27, CD69, and CD103 (**Fig. S1D**). P2RX7 expression on liver T-cell lineages also trended higher compared to their blood counterparts in unmatched blood donors, and also between one paired liver and blood sample (**Fig. 1A** and **S1D**). Overall, these data illustrate that steady-state expression of P2RX7 is a feature of human unconventional T-cell lineages.

### Mouse T-bet^+^ MAIT, γδT, and NKT cells highly co-express ARTC2 and P2RX7

We next used mouse models to examine the regulation of unconventional T cells by P2RX7 activation. Previous studies showed that expression of ARTC2 drives the NAD-mediated activation of P2RX7, and co-expression of ARTC2 and P2RX7 has been characterised on regulatory T cells and NKT-cell subsets (Di Virgilio et al., 2017; Rissiek et al., 2013; Schenk et al., 2011; Scheuplein et al., 2009; Seman et al., 2003). We analysed the expression of ARTC2 and P2RX7 on T cells across organs, focusing on the unconventional T-cell compartment and their functional signatures (**Fig. 1B-E** and **Fig. S2A-B**). For liver and spleen T cells, dimensionality reduction of flow cytometry analysis delineated clusters of MAIT, γδT, and NKT cells, defined by labelling with MR1-5-OP-RU tetramer, anti-γδTCR, and CD1d-α-GalCer tetramer, where the majority of MAIT and NKT cells, and a subset of γδT cells, expressed PLZF, the master transcriptional regulator of unconventional T cells, and CD44 (**Fig. 1C** and **Fig. S2A**). Co-expression of ARTC2 and P2RX7 on all T cells overlapped with markers such as CD38, CD44, CD69, and PLZF (**Fig. 1C, Fig. S2A**), indicative of effector-memory T cells that are tissue-associated (Stark et al., 2018). The MAIT, γδT, and NKT-cell clusters were divided into ARTC2^+^P2RX7^+^ and ARTC2^-^P2RX7^-^ populations, where this dichotomy appeared to correlate with T-bet^+^ and RORγt^+^ clusters, respectively (**Fig. 1C** and **Fig. S2A**). Accordingly, these data suggest that co-expression of ARTC2 and P2RX7 is a signature of multiple PLZF^+^ unconventional T-cell lineages in mice.

Across different organs, ARTC2^+^P2RX7^+^ unconventional T cells were significantly more abundant in liver and spleen than in thymus and inguinal lymph nodes (iLNs) (**Fig. 1B**). On average, 45%, 55%, and 90% of mouse liver MAIT, γδT, and NKT cells, respectively, were ARTC2^+^P2RX7^+^ (**Fig. 1B**). While most NKT cells in spleen co-expressed ARTC2 and P2RX7, only a minority of splenic MAIT and γδT cells were ARTC2^+^P2RX7^+^ (**Fig. 1B**). To determine whether ARTC2/P2RX7 expression was constrained to functionally distinct subsets of unconventional T cells, we examined them for transcription factor co-expression (**Fig. S2B**). This revealed that ARTC2^+^P2RX7^+^ cells in liver were primarily T-bet^+^ MAIT1, γδT1, and NKT1 cells (**Fig. 1D-E**). The results were similar in spleen, except for spleen γδT1 cells which were mostly P2RX7^-^. Furthermore, a subset of CD44^neg^ γδT cells co-expressed ARTC2 and P2RX7 in thymus, spleen, and iLNs (**Fig. S2C**). In contrast, few RORγt^+^ MAIT17, γδT17, and NKT17 cells were ARTC2^+^P2RX7^+^, except a subset of NKT17 cells from spleen and liver (**Fig. 1D-E**). Taken together, these data suggest that ARTC2 and P2RX7 are mainly co-expressed by T-bet^+^, but not RORγt^+^, unconventional T cells, with the highest expression detected in liver and, to a lesser extent, spleen.

Given that ARTC2 expression extended beyond MAIT and NKT cells in our analysis (**Fig. 1C**), we also examined non-MAIT/NKT αβT cells. This revealed that many (∼40-50%) CD44^hi^ CD4^+^ αβT cells co-expressed ARTC2 and P2RX7 in spleen and liver, and to a lesser extent, iLNs and thymus (**Fig. 1F-G** and **Fig. S2D**). A significantly greater proportion of CD44^hi^ CD8^+^ and CD44^hi^ CD4^-^CD8^-^ DN αβT cells were ARTC2^+^P2RX7^+^ in liver relative to spleen, iLNs, and thymus (**Fig. 1F-G** and **Fig. S2D**). A subset of liver CD44^hi^ CD4^+^ and CD44^hi^ CD4^-^CD8^-^ DN αβT cells expressed ARTC2 at a higher intensity (MFI) compared to ARTC2^+^P2RX7^+^ CD8^+^ T cells, though these ARTC2^hi^ αβT cells were much rarer or absent in spleen, iLNs, and thymus (**Fig. 1G** and **Fig. S2D-E**). In CD44^neg^ αβT cells in thymus, spleen, and liver, CD4^-^CD8^-^ DN αβT cells co-expressed intermediate levels of ARTC2 and P2RX7 (**Fig. 1F** and **Fig. S2D-E**). Lastly, most CD44^neg^ CD4^+^ and CD44^neg^ CD8^+^ αβT cells within peripheral tissues expressed ARTC2 at an intermediate level, though few expressed P2RX7 (**Fig. S2D-E**). As PLZF was only detected amongst CD44-expressing αβT and γδT cells (**Fig. 1H** and **Fig. S2F**), we next analysed the expression of PLZF within CD44^+^ ARTC2^hi^ non-MAIT/NKT αβT cells (**Fig. 1G** and **Fig. S2E**). Here, ARTC2^hi^ CD4^+^ and ARTC2^hi^ CD4^-^CD8^-^ DN αβT cells expressed PLZF and were predominantly T-bet^+^ (**Fig. 1H**). Out of all ARTC2^+^ γδT cells, only ARTC2^hi^ γδT cells expressed PLZF (**Fig. S2F**). Taken together, these data reveal populations of effector-like, PLZF^+^T-bet^+^ αβT cells that mainly reside in the liver alongside other ARTC2^hi^P2RX7^+^ unconventional T-cell types.

As P2RX7 may play a role in NKT-cell homeostasis (Bovens et al., 2020; Liu and Kim, 2019), we analysed the frequency and number of MAIT, γδT, and NKT cells, and non-MAIT/NKT αβT cells in *P2rx7*^-/-^ mice (**Fig. S3**). The absolute numbers of lymphocytes across organs were not significantly different between *P2rx7*^-/-^ and WT mice (**Fig. S3B, C**), except for in liver, where total lymphocytes were slightly higher in *P2rx7*^-/-^ mice relative to WT mice (**Fig. S3B**). All T-cell subsets were present at similar frequencies between *P2rx7*^-/-^ and WT mice, although some minor but statistically significant differences were detected (**Fig. S3C**). As the frequency of total ARTC2^+^ and ARTC2^hi^ T cells and their expression of ARTC2 was similar between *P2rx7*^-/-^ and WT mice, these data collectively suggest that P2RX7 does not affect ARTC2 expression by subsets of T cells (**Fig. S3C-D**) (Stark et al., 2018). We additionally examined the subset distribution of MAIT, γδT, and NKT cells in *P2rx7*^-/-^ mice. Percentages of spleen MAIT1, γδT1, and NKT1 cells and thymic MAIT1 cells were slightly but significantly decreased in *P2rx7*^-/-^ mice relative to WT mice, with a concomitant increase in spleen and thymus MAIT17-cell and splenic NKT17-cell frequencies (**Fig. S3E**). The numbers of liver MAIT1, γδT1, and NKT1 cells, and liver and spleen NKT17 cells were slightly increased in *P2rx7*^-/-^ mice relative to WT mice (**Fig. S3F**). These findings indicate that the expression of P2RX7 is not essential for the overall development of unconventional T-cell lineages apart from a minor influence on unconventional T-cell functional subset diversification in the thymus and spleen, and numbers in liver.

### ARTC2 and P2RX7 expression on T cells is induced by retinoic acid

The liver is a major storage site for retinol and participates in retinoic acid metabolism (Blaner et al., 2016), where the retinoic acid (RA)-induced, RA receptor alpha (RARα) can bind to the enhancer region of *P2rx7* (Hashimoto-Hill et al., 2017). RA can increase expression of both P2RX7 and ARTC2 on activated intestinal T-cell subsets, and P2RX7 on antigen-stimulated NKT cells (Hashimoto-Hill et al., 2017; Heiss et al., 2008; Liu and Kim, 2019). We thus tested whether exposure to RA can induce P2RX7 and ARTC2 on unconventional T cells without TCR stimulation, particularly on thymic subsets that have the lowest steady state expression of both molecules (**Fig. 2** and **Fig. 1D-E**). Thymocytes were enriched for mature T cells, including thymic MAIT, γδT, and NKT cells, which were confirmed to be predominantly ARTC2^-^P2RX7^-^ (**Fig. S4A**). These thymocytes were cultured in the presence of RA or vehicle control for 3 days (**Fig. 2A-E**). An increase in ARTC2 expression on cells following culture was observed within the vehicle control, which may be attributed to trace amounts of RA present in the fetal bovine serum used in the culture media (**Fig. 2B-C** and **Fig. S4A**) (Napoli, 1986). After RA treatment, a greater proportion (46%) of thymic PLZF^+^T-bet^+^ T cells expressed P2RX7 relative to vehicle-treated cells (8%), RA-treated PLZF^+^RORγt^+^ T cells (18%), and RA-treated PLZF^-^ T cells (2%) (**Fig. 2B**). In line with the expression of ARTC2 by mature thymocytes (Koch-Nolte et al., 1999), over half of thymic PLZF^-^ T cells were ARTC2^+^ after culture without RA (**Fig. 2B**). A moderate increase in the frequency of ARTC2^+^ cells was observed amongst thymic PLZF^+^T-bet^+^ and PLZF^-^ T cells, and to a lesser extent, thymic PLZF^+^RORγt^+^ T cells, after RA treatment (**Fig. 2B**).

**Figure 2:**
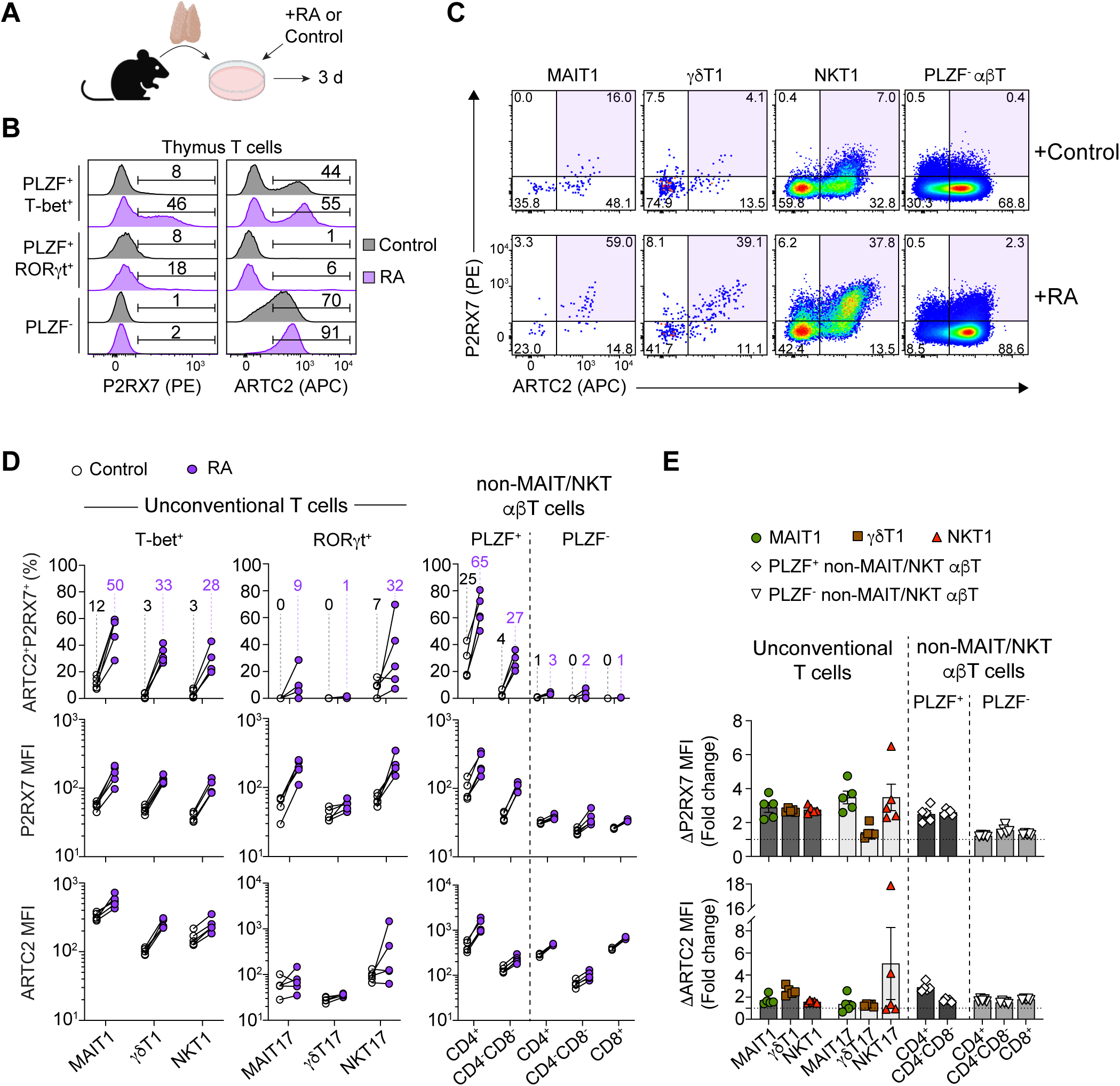
Retinoic acid induces expression of ARTC2 and P2RX7 on T cells. (A) Experimental schematic. Mouse thymocytes were cultured for three days in the presence of 20nM all-*trans* retinoic acid (RA) or the vehicle control. 3 – 5 thymuses were pooled prior to complemented-mediated depletion of immature CD24^+^ thymocytes. (B & C) Flow cytometric analysis of ARTC2 and P2RX7 expression by indicated thymic T-cell types 3 days after treatment with RA or the vehicle control. Histogram and FACS plots show concatenated data from all replicate samples from one of two similar experiments. (B) Numbers in histograms represent percentages of P2RX7^+^ and ARTC2^+^ cells of indicated T-cell types. (C) Numbers within FACS plots represent percentage of gated cells out of T-bet^+^ MAIT1, γδT1, and NKT1 cells, or PLZF^-^ non-MAIT/NKT αβT cells. (D) Graphs show the percentages of ARTC2^+^P2RX7^+^ cells of indicated T-cell types and mean fluorescence intensity (MFI) of P2RX7 and ARTC2 expression. White- and purple-shaded circles represent data from vehicle or RA-treated cells, respectively. Connecting lines represent paired data. Numbers above data points represent the average percentage for indicated data sets. (E) Graphs show fold change in P2RX7 and ARTC2 MFI amongst RA-treated cells relative to vehicle controls. Horizontal dotted lines represent a fold change of 1. (D & E) Each symbol represents data from 3 – 5 pooled thymuses where total of 5 pooled thymus samples were analysed across 2 separate experiments. (E) Graphs depict individual data points and mean ± SEM.

Within specific cellular lineages, a subset of thymic MAIT1, γδT1, and NKT1 cells, as well as PLZF^+^ non-MAIT/NKT αβT cells, upregulated both markers to become ARTC2^+^P2RX7^+^ after RA exposure compared to vehicle treatment (**Fig. 2C-E**). This was also reflected in the higher intensity in the expression of P2RX7 and ARTC2 (**Fig. 2C-E**). On average, an overall higher proportion of thymic MAIT1 (50%), γδT1 (33%), NKT1 (28%), and PLZF^+^ non-MAIT/NKT αβT cells (CD4^+^ = 65%, CD4^-^CD8^-^ = 27%) were ARTC2^+^P2RX7^+^ after RA treatment relative to MAIT17 (9%) and γδT17 (1%) cells, with the exception of NKT17 cells (32%) (**Fig. 2D**). Only a small subset of thymic PLZF^-^ αβT cells upregulated P2RX7 to become ARTC2^+^P2RX7^+^ in response to RA (**Fig. 2B-D**), where the increase in P2RX7 expression by PLZF^-^αβT cells was less pronounced compared to PLZF^+^ αβT cell types (**Fig. 2C-E**).

Though some peripheral MAIT, γδT, and NKT cells express ARTC2 and P2RX7 at steady state, the frequency of ARTC2^+^P2RX7^+^ MAIT1, γδT1, and NKT1 cells from spleen and liver was also increased after RA treatment (**Fig. S4C-D**). Increases in P2RX7 and ARTC2 expression occurred to varying extents amongst most T-cell types from the spleen and liver post-RA treatment, in line with previous reports activating peripheral T cells in the presence of RA *in vitro* (Hashimoto-Hill et al., 2017; Heiss et al., 2008; Liu and Kim, 2019) (**Fig. S4B-D**). These findings indicate that retinoic acid can induce the co-expression of ARTC2 and P2RX7 by T cells, and in particular thymic PLZF^+^T-bet^+^ unconventional T-cell types.

### Unconventional T cells undergo ARTC2/P2RX7-dependent cell death and loss of surface CD27

To investigate the effects of P2RX7 activation on unconventional T cells, we used the release of NAD by cells damaged during organ processing, which is sufficient to drive robust ARTC2-mediated ADP-ribosylation of P2RX7 *ex vivo* (Scheuplein et al., 2009; Seman et al., 2003). While ADP-ribosylation by ARTC2 can occur at 4°C, NAD-mediated P2RX7 activation only occurs at 37°C, leading to cell death and activating downstream ADAM metalloproteases that induce ectodomain shedding of CD27 from the cell surface (Johnsen et al., 2019; Moon et al., 2006). Thus, we analysed the *ex vivo* viability of MAIT, γδT, and NKT cells by labelling with Annexin V and 7-AAD. Compared to cells maintained at 4°C, a larger proportion of liver MAIT, γδT, and NKT cells were at early (Annexin V^+^ 7-AAD^-^) or late (7-AAD^+^) stages of cell death when incubated at 37°C compared to those maintained at 4°C (**Fig. 3A-B**). This increase in cell death was inhibited when cells were isolated from mice pre-treated with the anti-ARTC2 nanobody (NB) ‘s+16’ prior to organ harvest (Rissiek et al., 2013) or when they were incubated in the presence of the P2RX7 inhibitor, A438079 (**Fig. 3A-C**). Whilst increases in cell death were observed in spleen unconventional T cells at 37°C, only spleen NKT-cell death was rescued by A438079 treatment (**Fig. 3C**). As expected, no significant changes were seen in the proportion of 7-AAD^+^ MAIT, γδT, and NKT cells from thymus across all conditions (**Fig. S5A**).

**Figure 3:**
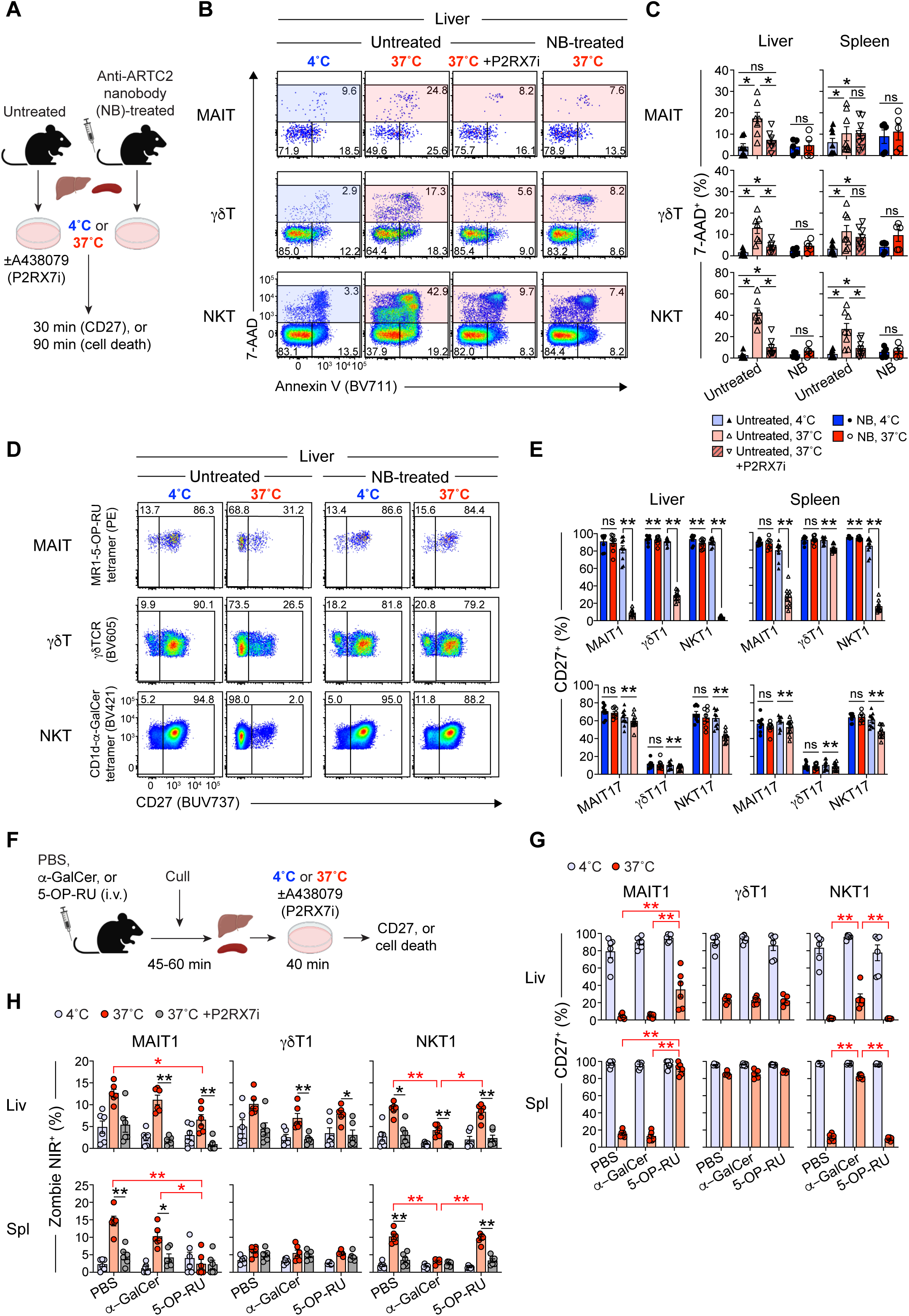
P2RX7 activation on unconventional T cells induces cell death and loss of surface CD27. (A) Experimental schematic. Liver and spleen cells from untreated or anti-ARTC2 nanobody (clone: s+16) (NB)-treated mice were cultured at 4°C or 37°C for 30 or 90 minutes prior to analysis. Untreated mouse cells were cultured with or without the P2RX7 inhibitor (P2RX7i), A438079 (10µM). (B) Flow cytometric analysis of liver MAIT, γδT, and NKT cells co-labelled with 7-AAD and Annexin V after a 90-minute incubation. (C) Graphs depict percentage of 7-AAD^+^ (dead) cells of total MAIT, γδT, and NKT cells from the liver and spleen. n = 3 separate experiments with a total of 4-8 mice/group. ns P>0.05, *P≤0.05 using a Mann-Whitney U test with correction for multiple comparisons for the untreated groups. (D) Flow cytometric analysis of surface CD27 expression on total liver MAIT, γδT, and NKT cells. (E) Graphs depict percentages of CD27^+^ cells out of viable T-bet^+^ MAIT1, γδT1, and NKT1 cells, and RORγt^+^ MAIT17, γδT17, and NKT17 cells. n = 3-4 separate experiments with a total of 8-10 mice/group. ns P>0.05, *P≤0.05, **P≤0.01 using a Wilcoxon matched pairs signed rank test. Numbers in FACS plots represent percentage of gated cells. (F) Experimental schematic. Mice were i.v. administered 5-OP-RU (200 pmol), α-GalCer (2µg), or PBS and liver and spleens were harvested 45-60 minutes later. Liver and spleen cells were cultured at 4°C or 37°C for 40 minutes with or without A438079 (10uM) prior to analysis. (G & H) Graphs depict percentages of viable CD27^+^ cells (G) or Zombie NIR^+^ (dead) cells (H) out of indicated cell-types. A total of 6 mice per group were analysed across 2 separate experiments. ns P>0.05 (not shown), *P≤0.05, **P≤0.01 using a Mann-Whitney U test with a Bonferroni-Dunn correction for multiple comparisons. (C, E, G, H) Graphs depict individual data points and mean ± SEM. Each symbol represents an individual mouse.

We also examined the loss of surface CD27 by MAIT, γδT, and NKT cells due to P2RX7 activation at 37°C. The percentages of CD27^+^ MAIT, γδT, and NKT cells, especially from liver and spleen, were markedly decreased relative to control cells maintained at 4°C (**Fig. 3D-E** and **Fig. S5B**). This was dependent on ARTC2 activity, as it was blocked by pre-treatment of mice with the anti-ARTC2 NB (**Fig. 3D-E**). Moreover, CD27 expression was also maintained on cells from *P2rx7^-/-^*mice incubated at 37°C (**Fig. S5C**). The loss of CD27 by liver and spleen MAIT, γδT, and NKT cells, and NKT cells from thymus and iLNs, was predominantly observed amongst T-bet^+^ subsets, with less or no impact on their RORγt^+^ counterparts (**Fig. 3E** and **Fig. S5D**). In addition to CD44^hi^ γδT cells (i.e., γδT1 cells), we observed a reduced frequency of CD27-expressing CD44^neg^ γδT cells and some non-MAIT/NKT cell αβT-cell subsets from liver, spleen, iLNs, and, to a lesser extent, thymus after incubation at 37°C (**Fig. S5E-F**). Expression of ARTC2 by liver MAIT, γδT, and NKT cells incubated at 37°C was lower in cells from untreated control mice compared to cells harvested from NB-treated mice (**Fig. S5G**), likely representing ARTC2 shedding following P2RX7 activation as previously reported on T cells (Menzel et al., 2015). Furthermore, P2RX7 was absent on liver MAIT, γδT, and NKT cells, regardless of temperature, without NB treatment, yet was clearly present on these cells when mice had been treated with the anti-ARTC2 NB, which is consistent with an earlier report on NKT cells (Borges Da Silva et al., 2019; Rissiek et al., 2013) (**Fig. S5G**). Together, these findings indicate that exposure to NAD released by damaged cells can result in the NICD of liver and spleen unconventional T cells in a manner dependent on ARTC2 activity and P2RX7 activation.

Given that TCR stimulation can modulate the sensitivity of some T cell subsets to ARTC2-mediated P2RX7 activation (Faliti et al., 2019; Kahl et al., 2000; Proietti et al., 2014; Stark et al., 2018), we investigated the loss of CD27 and death of T-bet^+^ MAIT1 and NKT1 cells upon cell culture following cognate antigen encounter *in vivo* (**Fig. 3F-H**) After intravenous administration of 5-OP-RU or α-GalCer, to stimulate MAIT and NKT cells, respectively, liver MAIT and NKT cells upregulated CD69, indicative of their activation *in vivo* (**Fig. S6B**). In contrast to PBS-treated mice, a significantly higher percentage of liver and spleen MAIT1 and NKT1, but not γδT1 cells, from 5-OP-RU and α-GalCer treated mice were CD27^+^ and a significantly lower percentage of these cells were undergoing cell death (Zombie NIR^+^) after the culture period (**Fig. 3G-H**). 5-OP-RU-exposed MAIT1 and α-GalCer-exposed NKT1 cells had greatly reduced ARTC2 expression (**Fig. S6C-D**), in line with ARTC2 shedding following TCR stimulation. Lastly, no decrease in P2RX7 expression was found on stimulated MAIT1 and NKT1 cells (**Fig. S6C-D**), in contrast to previous reports in CD8^+^ tissue-resident memory and T follicular helper T cells (Faliti et al., 2019; Proietti et al., 2014; Stark et al., 2018). Instead, an increase in P2RX7 expression relative to PBS control mouse cells was observed (**Fig. S6C-D**), likely representing the loss of ARTC2-mediated P2RX7 downregulation *ex vivo* (Borges Da Silva et al., 2019). Overall, these results suggest that cognate antigen encounter by liver MAIT1 and NKT1 cells reduces their susceptibility to extracellular NAD and NICD via the modulation of surface ARTC2 expression rather than downregulation of P2RX7.

### ARTC2 blockade improves recovery of unconventional T cells following adoptive transfer

As temperature is a factor in unconventional T cells undergoing NICD *ex vivo*, it was likely that this would influence survival of these cells upon adoptive transfer *in vivo* (**Fig. 4** and **Fig. S7**). Lymphocytes harvested from liver and spleen of anti-ARTC2 NB-treated and untreated WT C57BL/6 Ly5.1 mice were labelled with CTV and CFSE, respectively, mixed at a 1:1 ratio, and transferred into WT Ly5.2 recipients. Though both NB-treated (CTV^+^) and untreated (CFSE^+^) donor cells were recovered from liver 8 days post-transfer, there was a strong and significant bias toward NB-treated unconventional T cells, particularly NKT and MAIT cells, and to a lesser extent, γδT cells regardless of their liver (**Fig. 4B-C**) or spleen (**Fig. 4D-E**) origin. The bias towards NB-treated cells was also reflected in the ratios of NB-treated to untreated cells following adoptive transfer (**Fig. S7A**). In line with the comparatively low co-expression of ARTC2 and P2RX7 by splenic γδT cells, the average ratio of NB-treated to untreated splenic γδT cells recovered from the spleen was similar to that prior to transfer (**Fig. 1B, D, E** and **Fig. S2C**), As a control for the efficiency of adoptive transfer from the different donors, the collective population of non-T/non-B cells was analysed and were recovered at a ∼1:1 ratio of NB-treated to untreated cells (**Fig. 4B-E** and **Fig. S7A, B**), though there was a slight but significant increase in NB-treated, liver-derived non-T/non-B cells recovered from liver. In peripheral LNs (pLNs), a significant increase in NB-treated, liver-derived γδT cells was found, though this was not observed amongst MAIT cells or spleen-derived γδT cells (**Fig. S7B**).

**Figure 4.**
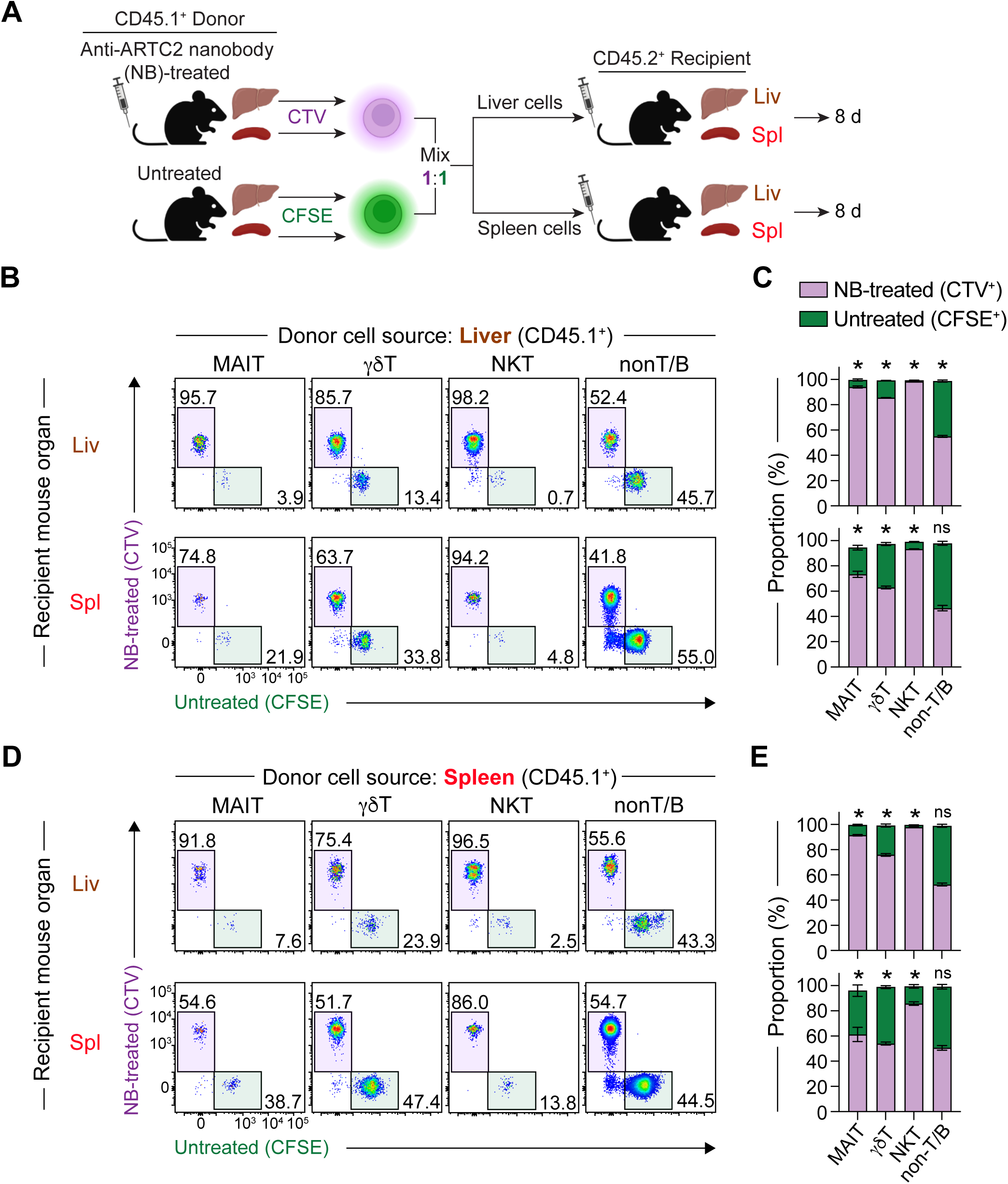
ARTC2 blockade improves recovery of adoptively transferred unconventional T cells. (A) Experimental schematic. Liver and spleen cells from anti-ARTC2 nanobody (clone: s+16) (NB)-treated or untreated mice were labelled with CTV (NB-treated) or CFSE (untreated) prior to co-transfer at a 1:1 ratio into recipient mice. Donor liver and spleen cells were recovered from recipient mouse livers and spleens 8 days later. Donor and recipient spleen cells were subjected to magnetic bead depletion of B220^+^ and CD62L^+^ cells, or B220^+^ cells, respectively. (B & C) Flow cytometric analysis of donor CD45.1^+^ MAIT, γδT, NKT, and non-T/B cells sourced from liver (B) or spleen (C) and recovered from liver and spleens of recipient mice 8 days after adoptive transfer. Numbers in FACS plots represent percentage of gated cells. MAIT and γδT-cell FACS plots were generated by concatenation of data from all (n = 3) mice of the same group from one of two similar experiments. (C & E) Stacked bar charts depict the percentages of donor CTV^+^ (purple) and CFSE^+^ (green) cells upon recovery from recipient mice on day 8. Graphs depict individual data points and mean ± SEM. n = 2 separate experiments with a total of 6 recipient mice per group. ns P>0.05, *P≤0.05, **P≤0.01 using a Wilcoxon matched-pairs signed-rank test for comparison between NB-treated vs untreated.

Consistent with the earlier findings that ARTC2 and P2RX7 were predominantly expressed by MAIT1, γδT1, and NKT1 cells, increases in donor cell recovery reflected significantly increased recovery of these subsets, as defined by surface surrogate markers CD44 and CD319 (CD44^+^CD319^+^) for these cells (Xu et al., 2023), although MAIT17 and NKT17 cells (defined as ICOS^+^CD319^-^), and γδT17 cells (CD44^hi^CD319^-^) were also increased in some cases (**Fig. S7C**). Accordingly, these data demonstrate that anti-ARTC2 blockade within donor mice markedly improves the survival and recovery of MAIT1, γδT1, and NKT1 cells after adoptive transfer, particularly for MAIT and NKT cells, and liver-derived donor T-cell subsets.

### P2RX7 activation primarily depletes T cells that co-produce IFN-γ and IL-4

Given the differential impact of ARTC2-mediated P2RX7 activation on T-bet^+^ and RORγt^+^ unconventional T cells, we examined its effect on cytokine production by these cells (**Fig. 5**). On average, 15%, 43%, and 46% of liver MAIT, CD44^+^ γδT, and NKT cells, respectively, produced IFN-γ after stimulation with PMA and ionomycin (**Fig. 5B-C**). Pre-treating mice with the anti-ARTC2 NB increased these percentages to, on average, 50%, 57%, and 94%, respectively (**Fig. 5B-C**). Similar increases were observed when cells from NB-untreated mice were stimulated in the presence of the P2RX7 inhibitor, A438079, or when comparing stimulated cells from WT and *P2rx7*^-/-^ mice (**Fig. 5B-C** and **Fig. S8A**). Smaller increases were found amongst IFN-γ-producing MAIT and NKT cells from spleen, and no increase was seen in the percentage of IFN-γ-producing γδT cells (**Fig. 5C** and **Fig. S8A-B**). Furthermore, the frequencies of IFN-γ-producing non-MAIT/NKT αβT cells from liver, but not spleen, were significantly increased when stimulated in the presence of A438079 (**Fig S8C**). In contrast, and as expected, similar percentages of IFN-γ-producing cells from thymus were observed across all conditions and between WT and *P2rx7*^-/-^ mice (**Fig S8A-C**). While significant reductions were seen in the percentages of IL-17-producing MAIT, γδT, NKT, and non-MAIT/NKT αβT cells following NB- or A438079-treatment, this likely reflected the increase in IFN-γ-producing cells with treatment (**Fig. 5B-C** and **Fig. S8B-C**), because the absolute numbers of IL-17-producing cells were largely similar across all conditions, with some minor exceptions (**Fig. 5C** and **Fig. S8C**). Accordingly, these data suggest that IFN-γ-but not IL-17-producing unconventional T-cell populations from liver are selectively regulated by ARTC2-mediated P2RX7 activation.

**Figure 5.**
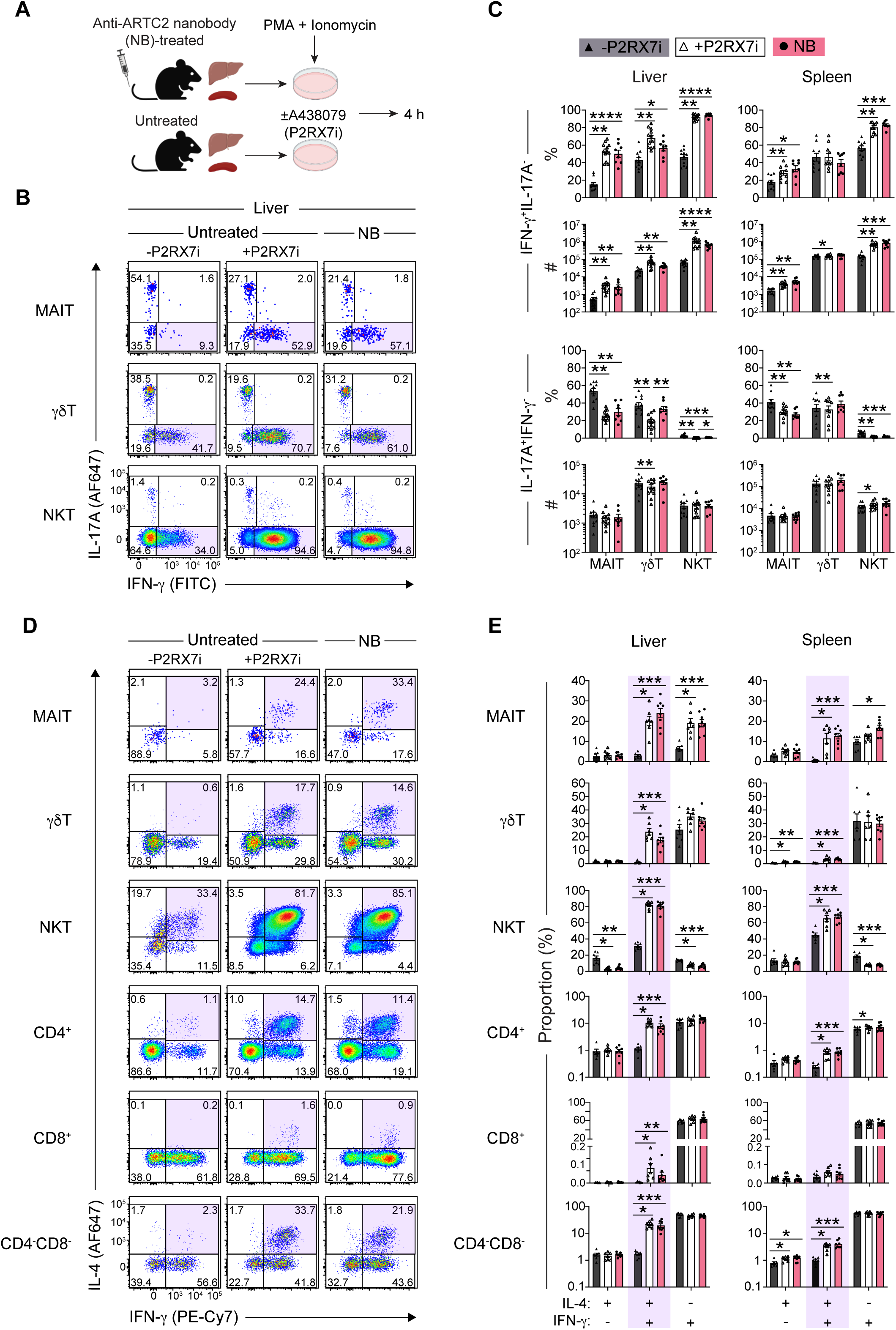
Blockade of ARTC2-mediated P2RX7 activation preserves unconventional T cells that co-produce IFN-γ and IL-4. (A) Experimental schematic. Liver and spleen cells from untreated or anti-ARTC2 nanobody (clone: s+16) (NB)-treated mice were stimulated with PMA and ionomycin and analysed 4 hours later. Untreated mouse cells were stimulated with or without the P2RX7 inhibitor (P2RX7i), A438079 (10µM). (B & D) Flow cytometric analysis of IL-17A and IFN-γ expression (B), and IL-4 and IFN-γ expression (D), by CD44^+^ MAIT, γδT, and NKT cells, and indicated CD44^+^ non-MAIT/NKT αβT-cell subsets. Numbers in FACS plots represent percentage of gated cells. (C & E) The percentage (%) and absolute number (#) of IL-17A^-^IFN-γ^+^ and IL-17A^+^IFN-γ^-^ MAIT, γδT, and NKT cells (C) and the percentages of IL-4/IFN-γ subsets amongst the specified T-cell subsets (E) were graphed. Each symbol represents an individual mouse. (C) n = 3-5 separate experiments with a total of 8-12 mice/group. (E) n = 3 separate experiments with a total of 7-8 mice/group. n.s. P>0.05 (not shown on graph) *P≤0.05, **P≤0.01, ***P≤0.001, and ****P≤0.0001 using a Wilcoxon matched-pairs signed rank test for +P2RX7i vs -P2RX7i or using a Mann-Whitney U test with a Bonferroni-Dunn correction for multiple comparisons for all other comparisons between conditions.

Whilst a subset of IFN-γ-producing NKT and γδT cells can simultaneously secrete IFN-γ and IL-4 (Azuara et al., 1997; Cameron and Godfrey, 2018; Gerber et al., 1999; Lee et al., 2013; Lee et al., 2015; Narayan et al., 2012; Pereira et al., 2013), a distinct population of IL-4-producing MAIT cells has remained elusive. We examined the impact of blocking P2RX7 activation on IL-4 production by MAIT, γδT, and NKT cells, as well as other αβT cells (**Fig. 5D-E** and **Fig. S8D**). In untreated WT mice, whilst IFN-γ^+^IL-4^+^ NKT cells represented on average 30% of liver NKT cells, and IFN-γ^+^IL-4^+^ MAIT and γδT cells were barely detectable in livers and spleens (**Fig. 5D-E**). However, with anti-ARTC2 NB- or A438079-treatment, around 20% of liver MAIT and γδT cells, and 80% of liver NKT cells, co-produced IFN-γ and IL-4 after stimulation, also reflected in significant increases in IFN-γ^+^IL-4^+^-cell numbers, respectively (**Fig. 5D-E** and **Fig. S8D**). Increases in the frequency and number of CD4^+^ and CD4^-^CD8^-^ DN non-MAIT/NKT αβT cells that co-produced IFN-γ and IL-4 were found in liver and, to a lesser extent, spleen, following P2RX7 inhibition or ARTC2 blockade (**Fig. 5D-E** and **Fig. S8D**). A small subset of IFN-γ^+^IL-4^+^ CD8^+^ T cells was also found in liver in these blockade experiments. Generally, the most prominent increases were seen in cells that produce both IFN-γ and IL-4 over cells that produce either cytokine alone, suggesting that P2RX7 activation impacts on the survival of the population of IFN-γ and IL-4 co-producing cells, rather than directly regulating IL-4 production (**Fig. 5D-E** and **Fig. S8D**). We also analysed cytokine production by various T-cell types from lungs, iLNs, and thymus following P2RX7 inhibition or NB-treatment (**Fig. S8D**). In these tissues, the number of IL-4- and/or IFN-γ-producing T-cell populations seemed less influenced by the treatment, with only some moderate increases observed in lungs after stimulation (**Fig. S8D**). Furthermore, IL-4 and/or IFN-γ-producing MAIT cells were rare within lungs, iLNs, and thymus, regardless of NB- or A438079-treatment (**Fig. S8D**). Accordingly, these data suggest that ARTC2-mediated P2RX7 activation predominantly targets T cells, including unconventional T cells, from liver and spleen and primarily affects subsets of these cells that co-produce IFN-γ and IL-4.

### Exogenous NAD selectively depletes liver T-bet^+^ PLZF^+^ARTC2^hi^ T cells in vivo

We next examined whether tissue-damaged associated release of metabolites directly depletes unconventional T cells *in vivo*, where administration of exogenous NAD is an established model of triggering P2RX7 within mouse models (Bovens et al., 2020; Kawamura et al., 2006; Liu and Kim, 2019; Stark et al., 2018). Thirty minutes after NAD administration, compared to PBS controls, there was a sharp decrease in the percentage of all PLZF^+^T-bet^+^ T cells in liver (**Fig. 6B**), including the frequency and number of T-bet^+^ MAIT1, γδT1, and NKT1 cells (**Fig. 6C-D**). This depletion was specific to T-bet^+^ cells in liver as the frequency and number of splenic T-bet^+^ MAIT1, γδT1, and NKT1 cells, and number of RORγt^+^ MAIT17, γδT17, and NKT17 cells in both spleen and liver were similar between NAD- and PBS-treated mice (**Fig. 6B-D** and **Fig. S9A**). Furthermore, PLZF^+^ARTC2^hi^ CD4^+^ and PLZF^+^ARTC2^hi^ CD4^-^CD8^-^ DN non-MAIT/NKT αβT cells from liver, but not spleen, were also markedly decreased in both frequency and number, in line with the overall loss of liver PLZF^+^ARTC2^hi^ T cells within NAD-treated mice (**Fig. 6B, D** and **Fig. S9A-C**). In contrast, the absolute numbers of non-PLZF^+^ARTC2^+^ cells in liver were unchanged between NAD- and PBS-treated mice (**Fig. S9C**). Aligning with the selective loss of ARTC2^hi^ T cells (**Fig. S9B-C**), pre-treatment of mice with the anti-ARTC2 NB prior to NAD administration at least partially blocked depletion of liver PLZF^+^T-bet^+^ARTC2^hi^ T cells, including MAIT1, γδT1, NKT1, and PLZF^+^ARTC2^hi^ non-MAIT/NKT αβT cells (**Fig. 6B-D** and **Fig. S9B-C**). This suggested that the NAD-induced depletion of these cells is, at least in part, ARTC2-dependent (**Fig. 6B-D** and **Fig. S9B-C**). In addition, we found that the MFI of T-bet, but not RORγt, within unconventional T-cell subsets from liver of NAD-treated mice was lower compared to corresponding cells from PBS control mice (**Fig. 6B-D** and **Fig. S9D**). This decrease in T-bet expression was seen to lesser extent in spleen unconventional T cells (**Fig. 6B** and **Fig. S9D**).

**Figure 6.**
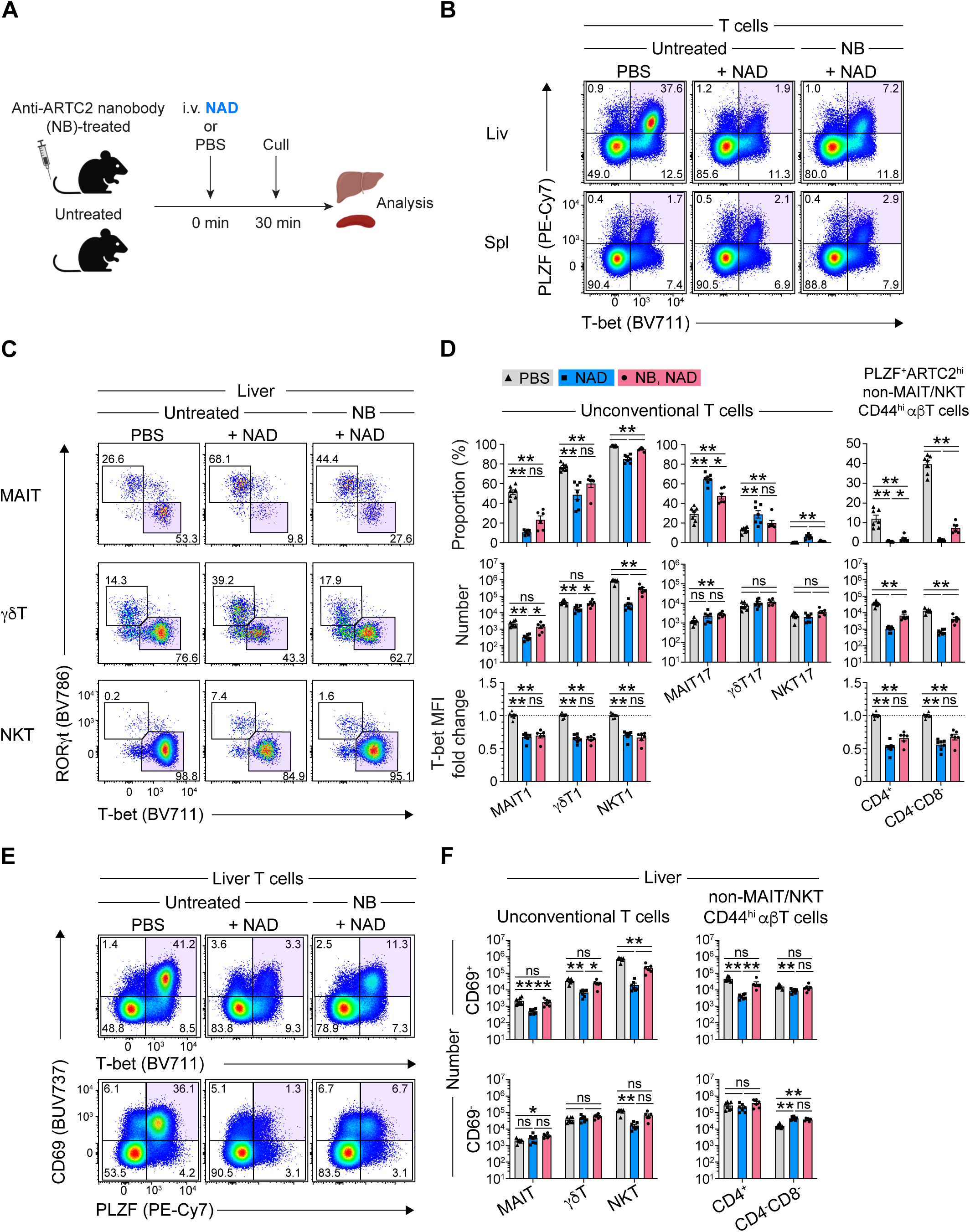
NAD selectively depletes liver T-bet^+^ unconventional T cells *in vivo*. (A) Experimental schematic. Anti-ARTC2 nanobody (clone: s+16) (NB)-treated or untreated mice were intravenously (i.v.) administered with nicotinamide adenine dinucleotide (NAD; 10mg) or PBS 30 minutes prior to organ harvest. (B & C) Flow cytometric analysis of PLZF and T-bet expression by liver T cells (B), and of RORγt and T-bet expression by liver MAIT, CD44^+^ γδT, and NKT cells (C). (D) Graphs depict the percentage and absolute number of, and fold change in the MFI of T-bet expression within the indicated liver T-cell types. Percentages of T-bet^+^ MAIT1, γδT1, and NKT1 cells, RORγt^+^ MAIT17, γδT17, and NKT17 cells are of total MAIT, γδT, and NKT cells. Percentages of PLZF^+^ARTC2^hi^ CD4^+^ and CD4^-^CD8^-^ cells are of total CD4^+^ and CD4^-^ CD8^-^ CD44^hi^ non-MAIT/NKT αβT cells. Fold change in T-bet expression is relative to PBS control mouse samples. (E) Flow cytometric analysis of CD69, T-bet, and PLZF expression, as indicated, by liver T cells. (F) Graphs depict the numbers of CD69^+^ and CD69^-^ MAIT, γδT, and NKT cells, and non-MAIT/NKT αβT-cell subsets from liver. n = 2 separate experiments with a total of 6-7 mice/group. Each symbol represents an individual mouse. n.s. P>0.05, *P≤0.05, **P≤0.01 using a Mann-Whitney U test with a Bonferroni-Dunn correction for multiple comparisons. (B, C, E) Numbers in FACS plots represent percentage of gated cells. (D & F) Graphs depict individual data points and mean ± SEM.

Given that exogenous NAD can deplete tissue-resident, CD69-expressing NKT and CD8^+^ memory T cells in liver (Bovens et al., 2020; Liu and Kim, 2019; Stark et al., 2018), we analysed whether other CD69^+^ unconventional T-cell subsets were affected by NAD administration (**Fig. 6E-F** and **Fig. S9E-G**). We found that CD69 marked most T-bet^+^, PLZF^+^, and ARTC2^hi^ T cells in livers of PBS-control mice (**Fig. 6E** and **Fig. S9E**). In contrast, only a subset of CD69^+^ T cells in spleen were ARTC2^hi^ and expressed T-bet or PLZF (**Fig. S9E-F**). After NAD administration, the frequency of total CD69^+^ T cells decreased in livers, but not spleens (**Fig. 6E-F** and **Fig. S9E-F**). The absolute numbers of liver CD69^+^ and CD69^-^ NKT cells were decreased in NAD-treated mice, although the decrease in the former subset was more pronounced (**Fig. 6F**), in line with a previous report (Bovens et al., 2020). CD69^+^, but not CD69^-^, MAIT, γδT, and non-MAIT/NKT CD44^hi^ αβT cells were decreased in livers of NAD-treated mice (**Fig. 6F** and **Fig. S9G**). Lastly, anti-ARTC2 NB pre-treatment of mice at least partially prevented the decrease in liver CD69^+^ cells (**Fig. 6F** and **Fig. S9G**). Accordingly, these data indicate that systemic exposure of mice to NAD selectively depletes multiple subsets of PLZF^+^T-bet^+^ unconventional T cells within liver.

## Discussion

Unconventional T cells, including MAIT, γδT, and NKT cells, and their functionally distinct subsets, are present in most tissues where lymphocytes are found, and are collectively abundant in liver (LeBlanc et al., 2022; Xu et al., 2023). How their rapid and diverse cytokine responses are regulated to balance immune defence while limiting inflammation is poorly understood. Here, we report that the tissue damage-associated metabolite NAD drives P2RX7 activation on multiple unconventional T-cell subsets, which directly impacts on their survival. In addition to NKT1 cells (Borges Da Silva et al., 2019), peripheral MAIT1 and γδT1 cells, along with a subset of CD4^+^ and CD4^-^CD8^-^ DN non-MAIT/NKT αβT cells, highly co-expressed ARTC2 and P2RX7 in liver, and to a lesser extent, spleen. Hepatic ARTC2^hi^ non-MAIT/NKT αβT cells were marked by expression of CD69, CD44, and P2RX7, as well as the transcription factors PLZF and T-bet, implying a similarity to other liver T-bet^+^ unconventional T-cell lineages, i.e. MAIT1, γδT1, and NKT1 cells. As T cells with intermediate expression of ARTC2 were PLZF^-^, this suggests that expression of PLZF may be linked to high expression of ARTC2 by T cells. It is plausible that factors within the tissue microenvironment may also drive high co-expression of ARTC2 and P2RX7 on PLZF^+^T-bet^+^ unconventional T cells. Supporting this, the liver is the main vitamin A storage site within the body (Blaner et al., 2016), where retinoic acid can induce expression of P2RX7 (*P2rx7*) and ARTC2 (*Art2b*) at both the protein and mRNA level on activated T cells (Hashimoto-Hill et al., 2017; Liu and Kim, 2019; Stark et al., 2018). Here, we show that retinoic acid upregulates the expression of P2RX7 and ARTC2 on PLZF^+^T-bet^+^ unconventional T cells, particularly those from thymus. Whilst the release of NAD by cells injured during routine organ processing (Scheuplein et al., 2009; Seman et al., 2003) was sufficient to drive the NICD of, and loss of surface CD27 expression by, unconventional T cells *ex vivo*, this was reduced amongst MAIT1 and NKT1 cells following cognate antigen encounter. Administration of exogenous NAD also rapidly depleted liver PLZF^+^ARTC2^hi^T-bet^+^ T cells, over other T-cell types, *in vivo*. In addition to P2RX7 inhibition reducing cell death *ex vivo*, nanobody-mediated ARTC2 blockade improved the recovery of adoptively transferred unconventional T cells and partly rescued their NAD-induced depletion *in vivo*, indicating a role for ARTC2-dependent P2RX7 activation in these cells’ survival. Due to rapid degradation of NAD *in vivo* (Adriouch et al., 2007; Cockayne et al., 1998), we cannot formally exclude the role for NAD-derived breakdown metabolites, such as ADP-ribose (Kawamura et al., 2006), in promoting the depletion of liver unconventional T cells *in vivo*.

Our findings suggest that blockade of ARTC2-dependent P2RX7 activation should be employed in studies of unconventional T cells, and indeed all peripheral T cells, from liver and spleen *ex vivo*. This will minimise alterations such as loss of surface CD27 expression and/or death of subsets of these cells, caused by exposure to NAD upon their isolation. Thus, key considerations for future studies, and interpretation of past studies include: i) the loss of surface CD27 by γδT1 cells upon cell culture, which may negate its use as a surrogate marker to identify IFN-γ-producing γδT cells (Ribot et al., 2009); ii) poor survival of T-bet^+^, IFN-γ and IL-4 co-producing T cells following isolation from tissues, which may explain the lower amounts of IFN-γ and IL-4 produced by peripheral MAIT, NKT, and γδT cells relative to their thymic counterparts after stimulation *ex vivo*; and iii) the previously unappreciated population of IFN-γ and IL-4 co-producing MAIT cells.

In line with this, a key finding in this study was the association between regulation by ARTC2-mediated P2RX7 activation, capacity to co-produce IFN-γ and IL-4, and co-expression of T-bet and PLZF. These findings support previous links between PLZF expression and the ability to co-produce IFN-γ and IL-4 within NKT and Vγ1^+^V86.3^+^ γδT cells (Kovalovsky et al., 2008; Kreslavsky et al., 2009), which is further corroborated by co-production of these cytokines by MAIT cells and subsets of non-MAIT/NKT CD4^+^ and CD4^-^CD8^-^ αβT cells. These observations are in line with the notion that PLZF expression imbues developing T cells with an effector-memory phenotype and ‘innate-like’ attributes (Kovalovsky et al., 2010; Kovalovsky et al., 2008; Kreslavsky et al., 2009; Pellicci et al., 2020; Savage et al., 2008). It will be intriguing to explore if the PLZF^+^ αβT cells examined here undergo a similar developmental program to that of MAIT, γδT, and NKT cells in acquiring PLZF expression intrathymically. Though IFN-γ and IL-4 are often considered to mediate functionally opposing immune responses, the role for PLZF^+^ unconventional T cells that can rapidly produce both cytokines in these immune contexts remains unclear. As cells that produced IL-17 alone were less susceptible to death by ARTC2-mediated P2RX7 activation, this suggests that this pathway may act to primarily control innate-like T cells that produce IFN-γ and/or IL-4, in particular those found within the liver and spleen.

Furthermore, IFN-γ and IL-4 both have pleiotropic effects within and outside of the immune system, and thus it is tempting to postulate that tissue damage-induced P2RX7 activation acts to prevent inappropriate and/or excessive cytokine production by peripheral unconventional T cells, such as in the context of sterile liver injury (Woolbright and Jaeschke, 2017). Several studies performed on individual T-cell lineages inform this speculation. For example, signalling through CD27 is known to promotes the survival, expansion, and function of γδT1 cells (Ribot et al., 2010; Ribot et al., 2009), suggesting that the ARTC2/P2RX7-dependent ectodomain shedding of CD27 from the surface of γδT1 cells may also act to supress IFN-γ production by these cells. In addition, IL-12, a pro-inflammatory cytokine which can stimulate unconventional T cells to produce IFN-γ in a TCR-independent manner (Darrigues et al., 2022), upregulates P2RX7 expression on CD8^+^ T cells (Stark et al., 2018). Though it is unknown whether IL-12 has a similar effect on unconventional T cells, perhaps heightened P2RX7 expression upon inflammation may sensitize cells to NICD. In turn, this axis may prevent the over-activation of IFN-γ-producing T cells in the context of inflammation, thereby reducing tissue damage and immunopathology.

It has been shown that ARTC2 can be shed from the surface of T cells after their activation (Kahl et al., 2000) or following P2RX7 activation (Menzel et al., 2015). ARTC2 in solution can then ADP-ribosylate cytokines such as IFN-γ, interfering with its ability to signal through IFN-γ receptors (Menzel et al., 2021). Accordingly, liver ARTC2^hi^P2RX7^+^ T cells may represent a reservoir of ARTC2 in a membrane bound state (Menzel et al., 2021), where upon T-cell activation and/or tissue damage, the release of ARTC2 may function to limit IFN-γ-mediated hepatic immune responses. This is supported by the rapid loss of ARTC2 by activated MAIT1 and NKT1 cells in liver and spleen following antigen-encounter *in vivo*, which was associated with their increased resistance to CD27 loss and cell death *ex vivo.* These findings suggest that TCR stimulation can rescue PLZF^+^T-bet^+^ unconventional T cells from tissue damage-induced, ARTC2/P2RX7-mediated cell death, supporting the notion that recently activated T cells are resistant to NICD (Rissiek et al., 2014; Rivas-Yáñez et al., 2020). The ARTC2-P2RX7 axis and its modulation following TCR signalling may act to promote the survival of PLZF^+^T-bet^+^ unconventional T cells actively responding to infection over bystander cells inadvertently activated by inflammatory stimuli, aligning with similar notions in T follicular helper cells (Faliti et al., 2019; Proietti et al., 2014) and CD8^+^ tissue-resident memory T cells (Stark et al., 2018).

As the ARTC2 gene is nonfunctional in humans (Haag et al., 1994), it will be important for future studies to elucidate whether P2RX7 activation in the collective human unconventional T-cell lineages are similarly subjected to ADP-ribosylation by orthologous ADP-ribosyltransferases in response to distinct tissue damage signals. Plausible family members within the human ARTC genes, such as ARTC1 and ARTC5, contain catalytic motifs for NAD binding and may play a similar role to mouse ARTC2 in terms of ADP-ribosylation of P2RX7 (Laing et al., 2011; Leutert et al., 2018)Cortés-Garcia et al., 2016; Hesse et al., 2022; Wennerberg et al., 2022). Although human unconventional T cells expressed P2RX7 to a greater extent than conventional T cells, we did not observe evidence for their NICD *in vitro*, in line with a previous report on human regulatory T cells (Cortés-Garcia et al., 2016). However, the overexpression of ARTC1 within some human cancers relative to normal tissues (Lin et al., 2024; Tang et al., 2013; Wennerberg et al., 2022) may drive the NICD of unconventional T cells within the tumours. Overall, there is emerging evidence describing an interplay where phenotypically- and functionally-synonymous subsets of unconventional T-cell lineages can exist within a competitive or compensatory niche, highlighting that they may be regulated by overlapping homeostatic and/or environmental cues (Ataide et al., 2022; Xu et al., 2023).

In summary, this study thus identifies the ARTC2-P2RX7 axis as a common modulator of PLZF^+^T-bet^+^ T-cell survival in the presence of tissue damage and damage-associated metabolites. This has a profound effect on unconventional T-cell lineages including MAIT, γδT, and NKT cells, but it is also more broadly active on other PLZF-expressing CD4^+^ and CD4^-^CD8^-^ αβT cells. These findings pose important questions for the collective regulation of unconventional T cells and the role for the ARTC2-P2RX7 axis in various disease contexts, also highlighting the need to inhibit this axis for *ex vivo* studies of liver and spleen T cells.

## Methods

### Mice

6- to 12-week-old C57BL/6 (B6) WT mice were bred in-house in the Department of Microbiology and Immunology Animal House, University of Melbourne. 10- to 12-week-old C57BL/6 Ly5.1 mice were purchased from ARC. *P2rx7*^-/-^ mice (Solle et al., 2001) and matched B6 WT controls were bred in-house at the Department of Anatomy and Physiology Animal House, University of Melbourne . In some of these experiments, as indicated, additional B6 WT controls came from the Department of Microbiology and Immunology Animal House, University of Melbourne. All mice used in experiments were housed under specific-pathogen-free conditions and were age- and sex-matched. All procedures on mice were approved by the University of Melbourne Animal Ethics Committee (#1914739, #21651, and #24324).

### Anti-ARTC2.2 nanobody administration and P2RX7 inhibition

Where specified, mice injected intravenously (i.v.) injected with 50µg of the anti-ARTC2 blocking nanobody (NB) (s+16a; Treg Protector, BioLegend) diluted in PBS 30 minutes prior to culling and organ harvest, as previously reported (Rissiek et al., 2013). The P2RX7 inhibitor, A438079 hydrochloride (A438079), (Santa Cruz Biotechnology) was dissolved in DMSO to a final concentration of 50mM. To prevent activation of P2RX7, A438079 was added to cell suspensions at 4°C and prior to incubation at 37°C, or as otherwise indicated.

### Preparation of cell suspensions

#### Mouse

Thymus, spleen, and lymph node cell suspensions were processed via gentle grinding through a 30µM nylon cell strainer into ice-cold FACS buffer (PBS with 2% FCS). Spleen cell suspensions were treated with red blood cell lysis buffer (Sigma-Aldrich) for 5 minutes at room temperature before washing with FACS buffer. Liver cell suspensions were prepared by gently grinding liver tissue through a 70µM nylon cell strainer into ice-cold FACS buffer. Liver leukocytes were isolated via density gradient centrifugation using a 33% Percoll (Cytiva) solution at room temperature. Liver cells were subsequently treated with red blood cell lysis buffer for 10 minutes at room temperature before washing with FACS buffer. In some experiments, thymus cell suspensions pooled from 3-5 mice were labelled with anti-CD24 (J11D) prior to complement-mediated (GTI Diagnostics) depletion of immature CD24^+^ cells to enrich for mature T cells. After depletion, viable CD24^-^ cells were isolated via density gradient centrifugation using Histopaque-1083 (Sigma-Aldrich). Where specified, spleen cell suspensions were subjected to a Histopaque-1083 density gradient to isolate viable cells. Prior to harvest, lungs tissues were perfused with PBS. Lungs were minced into small pieces and digested enzymatically in collagenase type III (Worthington Biochemical Corporation; 3mg/mL in RMPI-1640 supplemented with 2% FCS) in the presence of DNase I (Roche; 5µg/mL) for 90 minutes at 37°C, with or without the P2RX7 inhibitor as specified. After digestion, cell suspensions were treated with red blood cell lysis buffer (Sigma-Aldrich) for 5 minutes at room temperature before washing with FACS buffer. All cell suspensions were kept on ice or at 4°C unless otherwise specified.

#### Human

Peripheral blood mononuclear cells (PBMCs) were isolated from healthy human peripheral blood donors via standard density gradient centrifugation using Ficoll-Paque Plus (Cytiva). PBMCs were either used in experiments on the same day as processing or cryopreserved in liquid nitrogen for use at a later date. Human liver cell suspensions were generated either by gentle grinding tissue pieces through a 70µM nylon cell strainer or via mechanical separation using a gentleMACS Dissociator (Miltenyi Biotec). Liver cell suspensions were subjected to density gradient centrifugation using 33% Percoll prior to treatment with red blood lysis buffer. All liver cell suspensions were cryopreserved in liquid nitrogen prior to analysis. All procedures on human samples were approved by the University of Melbourne Ethics Committee (#2023-13000-42773-7 and human tissue immune responses #13009), and ADTB (HREC/48184/Austin-2019).

#### Tetramer Assembly

As previously described (Corbett et al., 2014; Reantragoon et al., 2013), tetramers of human and mouse MR1-5-OP-RU were generated in-house by refolding MR1 monomers in the presence of 5-A-RU and methylglyoxal. Enzymatic biotinylation of MR1-5-OP-RU monomers was conducted in the presence of the BirA enzyme. Biotinylated monomers were purified through size exclusion chromatography. Soluble monomers of human and mouse CD1d/μ_2_m protein were made in-house and biotinylated as previously reported (Gherardin et al., 2018). In brief, human and mouse CD1d-α-GalCer monomers were made by loading purified CD1d-biotin protein with the α-GalCer analogue, PBS-44, a gift from P. Savage (Brigham Young University, Provo, UT). This was conducted at a 6:1 (lipid:protein) molar ratio at room temperature overnight. Biotinylated monomers of MR1-5-OP-RU were tetramerised to streptavidin conjugated to PE (SAV-PE; Invitrogen) or BV421 (SAV-BV421; BioLegend). Biotinylated monomers of MR1-5-OP-RU or CD1d-α-GalCer were tetramerised to streptavidin conjugated to PE (SAV-PE; BD Pharmingen), BV421 (SAV-BV421; BioLegend), or BV785 (SAV-BV785; BioLegend). All streptavidin conjugates were added to biotinylated monomers across a series of 5 additions of one-fifth of the required volume separated by 8-10 minute incubations at 4°C and in the dark, and with immediate mixing after each addition.

### Flow cytometry

#### Surface staining

Cells were labelled with 7-aminoactinomycin D (7-AAD; Sigma-Aldrich) or the Zombie NIR fixable viability dye (BioLegend) and MR1-5-OP-RU tetramers conjugated to PE (Invitrogen) or BV421 (BD Biosciences) for 30 minutes at 4°C or at room temperature and in the dark. After washing, cells were labelled with CD1d-α-GalCer tetramers conjugated to BV421, PE, or BV785 alongside a panel of cell-surface monoclonal antibodies (Table 1) for 30 minutes at 4°C or at room temperature and in the dark.

**Table 1.** List of monoclonal antibodies and live/dead cellular dyes used.

| Used in Mouse Experiments |  |  |  |
| --- | --- | --- | --- |
| Specificity | Fluorochrome | Clone | Company/Source |
| CD16/32 | N/A | 2.4G2 | Produced in-house |
| 7-AAD | N/A | N/A | Sigma-Aldrich |
| Zombie NIR Fixable Viability Dye | N/A | N/A | BioLegend |
| Annexin V | BV711 | N/A | BD Horizon |
| Annexin V | FITC | N/A | BD Pharmingen |
| ARTC2 | APC | Nika102 | Novus Biologicals |
| B220 (CD45R) | BUV496 | RA3-6B2 | BD Horizon |
| B220 (CD45R) | BV786 | RA3-6B2 | BD Horizon |
| CD3 | BUV395 | 145-2C11 | BD Horizon |
| CD4 | BUV395 | GK1.5 | BD Horizon |
| CD4 | AF532 | RM4-5 | Invitrogen |
| CD8α | BUV805 | 53-6.7 | BD Horizon |
| CD27 | BUV737 | LG.3A10 | BD Horizon |
| CD27 | BV605 | LG.3A10 | BD Horizon |
| CD38 | FITC | 90/CD38 | BD Pharmingen |
| CD38 | BV711 | 90/CD38 | BD OptiBuild |
| CD44 | AF700 | IM7 | Invitrogen |
| CD45.1 | PE-Cy7 | A20 | BD Pharmingen |
| CD62L | APC | MEL-14 | Invitrogen |
| CD62L | PerCP-Cy5.5 | MEL-14 | Invitrogen |
| CD69 | BUV737 | H1.2F3 | BD Horizon |
| CD319 | APC | 4G2 | BioLegend |
| ICOS (CD278) | BV711 | C398.4A | BioLegend |
| IFN-γ | PE-Cy7 | XMG1.2 | BioLegend |
| IFN-γ | FITC | XMG1.2 | BD Pharmingen |
| IL-4 | AF647 | 11B11 | BD Pharmingen |
| IL-17A | AF647 | TC11-18H10 | BD Pharmingen |
| IL-17A | PerCP-Cy5.5 | TC11-18H10 | BD Pharmingen |
| P2RX7 | PE | 1F11 | BioLegend |
| P2RX7 | PE-Cy7 | 1F11 | BioLegend |
| PLZF | PE-Cy7 | 9E12 | BioLegend |
| RORγt | BV786 | Q31-378 | BD Horizon |
| RORγt | PerCP-Cy5.5 | Q31-378 | BD Pharmingen |
| T-bet | AF488 | 4B10 | BioLegend |
| T-bet | BV711 | 4B10 | BioLegend |
| TCRβ | APC-Cy7 | H57-597 | BD Pharmingen |
| $\gamma\delta$TCR | BV605 | GL3 | BioLegend |
| $\gamma\delta$TCR | BV605 | GL3 | BD OptiBuild |
| $\gamma\delta$TCR | FITC | GL3 | BD Pharmingen |
| **Used In Human Experiments:** |  |  |  |
| **Specificity** | **Fluorochrome** | **Clone** | **Company/Source** |
| 7-AAD | N/A | N/A | Sigma-Aldrich |
| LIVE/DEAD Blue<br>Near-IR | N/A | N/A | Invitrogen |
| LIVE/DEAD Fixable<br>Near-IR | N/A | N/A | Invitrogen |
| Zombie NIR Fixable<br>Viability Dye | N/A | N/A | BioLegend |
| CD3 | AF700 | UCHT2 | BD Pharmingen |
| CD3 | BUV395 | UCHT2 | BD Horizon |
| CD4 | BUV496 | SK3 | BD Horizon |
| CD4 | AF532 | RM4-5 | Invitrogen |
| CD8α | BUV805 | SK1 | BD Pharmingen |
| CD14 | BUV395 | MϕP9 | BD Pharmingen |
| CD14 | APC-Cy7 | MϕP9 | BD Pharmingen |
| CD14 | BV570 | M5E2 | BioLegend |
| CD19 | APC-Cy7 | SJ25C1 | BD Pharmingen |
| CD19 | BV570 | HIB19 | BD Pharmingen |
| CD19 | BUV737 | SJ25C1 | BioLegend |
| CD27 | BV785 | O323 | BioLegend |
| CD45 | AF700 | 2D1 | BioLegend |
| CD62L | FITC | DREG-56 | BD Pharmingen |
| CD161 | PerCP-Cy5.5 | HP-3G10 | BioLegend |
| CD161 | PE-Cy7 | HP-3G10 | BioLegend |
| FcR Blocking Reagent | N/A | N/A | Miltenyi Biotec |
| P2RX7 | AF647 | L4 | (Buell et al., 1998; Elhage et al., 2022)<br>Provided by Dr Xin Huang and Dr Ben Gu,<br>The University of Melbourne, Florey Institute<br>of Neuroscience and Mental Health. |
| Vα7.2 | BV711 | 3C10 | BioLegend |
| Vα7.2 | BV605 | 3C10 | BioLegend |
| Vδ1 | FITC | TS8.2 | Invitrogen |
| Vδ1 | VioBlue | REA173 | Miltenyi Biotec |
| Vδ2 | BV711 | B6 | BioLegend |
| Vδ2 | BUV563 | B6 | BD OptiBuild |
| $\gamma\delta$TCR | PE-Cy7 | 11F2 | BD |
| $\gamma\delta$TCR | R718 | 11F2 | BD OptiBuild |

#### Intracellular Transcription Factor Staining

After surface staining, cells were fixed and permeabilised using the eBioscience Foxp3/Transcription Factor staining kit (ThermoFisher Scientific) as per manufacturer guidelines. Cells were then stained for intranuclear transcription factors (Table 1) for 60-90 minutes at 4°C and in the dark prior to acquisition on a flow cytometer.

#### Intracellular Cytokine Staining

Cell suspensions generated from anti-ARTC2 NB-treated or untreated mice were stimulated with PMA (10ng/mL; Sigma-Aldrich) and ionomycin (1µg/mL; Sigma-Aldrich) in the presence of both GolgiStop (1:1500) and GolgiPlug (1:1000) (BD Biosciences) for 4 hours at 37°C. Cell suspensions from untreated mice were stimulated with or without the P2RX7 inhibitor, A438709 (10µM). A438709 was added to cold stimulation media before incubation at 37°C. After stimulation, cells were washed with FACS buffer and then stained for cell surface markers. Cells were then fixed and permeabilised in accordance with the manufacture’s guidelines using the BD Cytofix/Cytoperm kit (BD Biosciences). Cells were subsequently stained for anti-IL-17A (AF647 or PerCP-Cy5.5; clone TC11-18H10; BD Biosciences), anti-IFN-γ (FITC; clone XMG12; BioLegend, or PE-Cy7; clone XMG12; BD Biosciences), and anti-IL-4 (AF647; clone 11B11; BD Biosciences).

After labelling, cells were filtered through a 50µM mesh filter immediately before acquisition on a 5-laser (355 nm, 405nm, 488nm, 561nm and 633nm) BD LSR Fortessa or Cytek Aurora. Flow cytometric data was analysed using the FlowJo (BD) and OMIQ (Dotmatics) software.

### In vitro culture experiments

To analyse the loss of CD27 from the cell surface, mouse cell suspensions were incubated at 4°C or 37°C for 30 minutes before labelling with anti-CD27, unless specified otherwise. To analyse phosphatidylserine exposure, cell suspensions were incubated at 4°C or 37°C for 90 minutes and labelled with Annexin V-FITC or Annexin V-BV711 (BD Biosciences) in line with the manufacturer’s guidelines. In antigen stimulation experiments, where specified, mice were i.v. injected with 5-OP-RU (200 picomoles), α-GalCer (2µg), or PBS and culled 45-60 minutes later. Liver and spleen cell suspensions were incubated for 40 minutes with or without the P2RX7 inhibitor, A438079 (10µM), at 37°C prior to analysis of CD27 expression and cell death.

For cell culture experiments using RA (all-*trans* retinoic acid; Sigma, R2625), RA was first dissolved at 5mM in ethanol. Pooled thymuses from 3-5 mice were subjected to complement-mediated depletion of immature CD24^+^ thymocytes prior to cell culture. To facilitate survival of spleen and liver unconventional T cells *ex vivo*, mice were treated with the anti-ARTC2 NB prior to organ harvest. Prior to cell culture, viable spleen cells were isolated via density gradient centrifugation using Histopaque-1083. Cells were cultured with 20nM RA or with similarly-diluted vehicle control for 3 days prior to analysis.

For experiments involving cell culture of human PBMCs with exogenous NAD or ATP, NAD and ATP (Sigma) were dissolved in sterile PBS and pH was adjusted to ∼7.4. 100mM solutions of ATP and NAD were stored at -80°C as single-use aliquots. Freshly-isolated PBMCs were incubated for 10 minutes at 4°C with or without the P2RX7 inhibitor, A438079 (30uM) prior to incubation with NAD (0mM, 0.3mM, or 3mM) or ATP (3mM) for 2 or 18 hours at 37°C. Alternatively, cryopreserved human PBMCs were thawed using cell culture media pre-warmed to 37°C. Thawed PBMCs were incubated for 10 minutes at 37°C with or without A438079 (30µM) prior to culture with 3mM NAD or ATP for 18 hours, as indicated, in the presence of rhuIL-2 (50ng/mL; PeproTech). Within these experiments, mouse spleen cell suspensions were also incubated at 4°C for 10 minutes with or without the P2RX7 inhibitor, A438079 (30uM) prior to incubation with NAD (0.3mM or 3mM) for 2 hours at 37°C.

### Adoptive Transfer of cells

C57BL/6 WT Ly5.1 mouse donors were either treated with the anti-ARTC2 NB or left untreated 30 minutes before organ harvest. Liver and spleen cell suspensions were generated as above. To enrich for CD44^+^ MAIT, γδT, and NKT cells, spleen cells were depleted of B220- and CD62L-expressing cells via magnetic bead-based depletion (Miltenyi Biotec). Cells from anti-ARTC2 NB-treated or untreated donor mice were labelled with CTV (Invitrogen) or CFSE (Invitrogen), respectively, as per manufacturer’s guidelines. After labelling, donor cells were co-transferred at a 1:1 ratio of total CTV^+^-to-CFSE^+^ cells into C57BL/6 WT Ly5.2 recipients via an i.v. injection into the lateral tail vein. Recipients received either co-transferred liver or spleen cells. Donor cells were then recovered from recipient mice 8 days later from the liver, spleen, and pooled peripheral (inguinal, axillary, brachial) lymph nodes. To facilitate recovery of donor T cells from the spleen, recipient spleens were depleted of B220^+^ cells using magnetic beads as described above.

### NAD Administration

10mg of NAD (Sigma-Aldrich) was dissolved in PBS and pH was adjusted to ∼7.4 prior to i.v. injection into mice via the lateral tail vein. Mice were sacrificed 30 minutes later, and organs were collected for flow cytometric analysis.

## Funding and Acknowledgements

This work was supported by National Health and Medical Research Council of Australia (NHMRC) (1140126 and 2008913). H.F.K. is supported by an ARC DECRA (DE220100830), D.I.G. was supported by a Senior Research Fellowship (1117766) and subsequently by an NHMRC Investigator Grant (2008913), C.X. is supported by an Australian Postgraduate Award, and L.K.M is supported by the Sylvia and Charles Viertel Charitable Foundation. We gratefully acknowledge the generosity of the deceased organ donors and their families in providing valuable tissue samples to advance medical research. The Australian Donation and Transplantation Biobank (ADTB) is supported by the Australian Centre for Transplant Excellence and Research (ACTER), Austin Health Research Foundation and the Department of Microbiology and Immunology, Peter Doherty Institute for Infection and Immunity, University of Melbourne. The authors thank Garth Cameron for helpful discussions on the topic. We thank the staff in the Doherty Institute Animal House for animal husbandry assistance. We also thank Alexis Gonzalez, Vanta Jameson, and staff at the Melbourne Cytometry Platform (MCP) for flow cytometry support. All experimental schematics were created with BioRender.com.

## Author Contributions

C.X. and H.F.K. designed and performed experiments, analysed and interpreted results, and wrote the manuscript with input from D.I.G. A.O. and M.Q. assisted in experiments. A.F., G.S., R.D. and C.L.G coordinate organ donor sample collection through the ADTB. A.B., J.W., X.H., L.B., L.K.M., provided key reagents, mice, and intellectual input. D.I.G., and H.F.K. conceived and led the study. All authors approved the manuscript.

