## Supplementary Figures 1-9 for "Selective regulation of IFN-γ and IL-4 co-producing unconventional T cells by purinergic signalling"

Supp Fig. 1

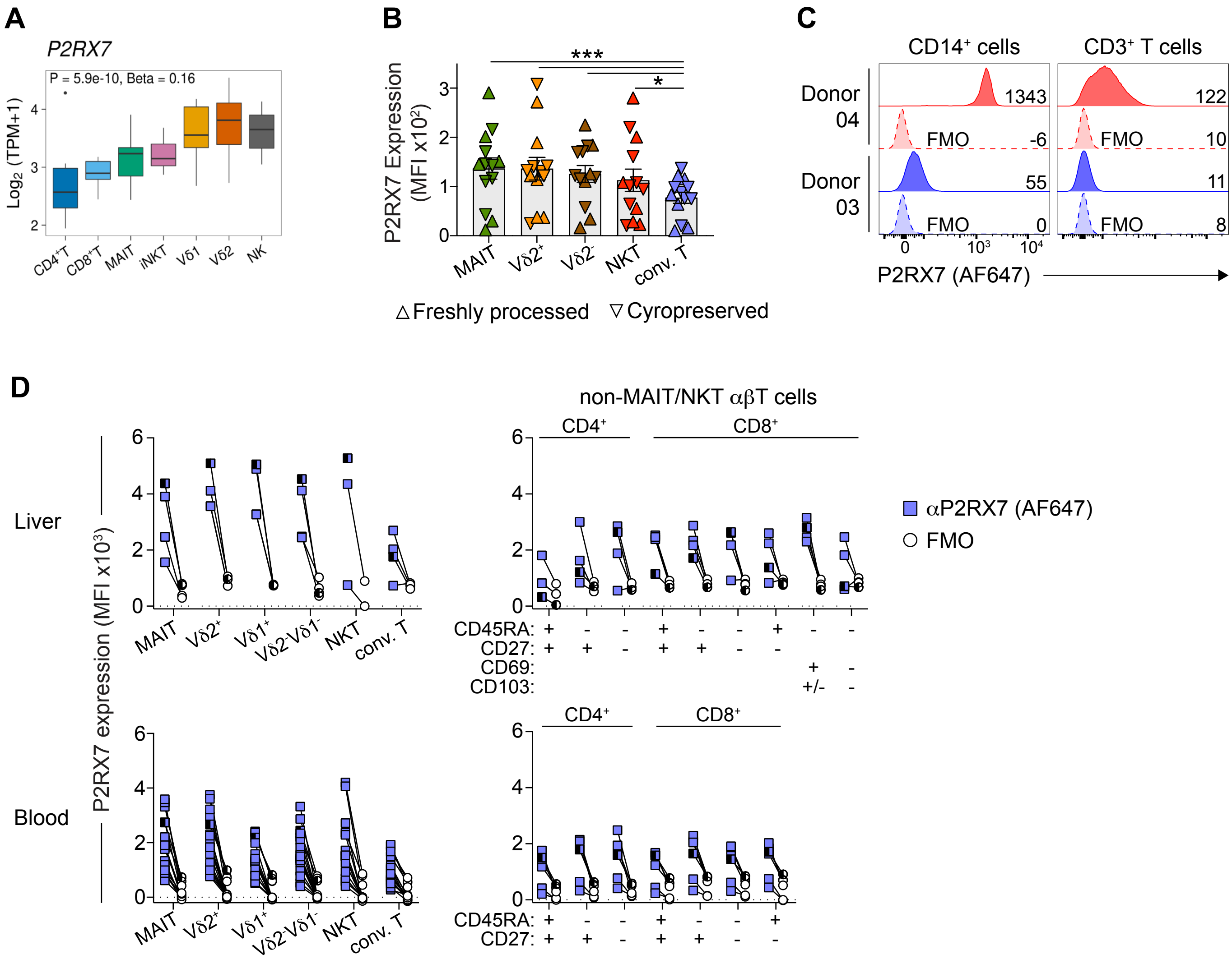
A
P2RX7
P = 5.9e-10, Beta = 0.16
Log2 (TPM+1)
CD4+T CD8+T MAIT iNKT Vδ1 Vδ2 NK
B
***
*
P2RX7 Expression (MFI x10²)
MAIT Vδ2+ Vδ2- NKT conv. T
△ Freshly processed ▽ Cyropreserved
C
CD14+ cells
CD3+ T cells
Donor 04
1343
122
FMO
-6
10
Donor 03
55
11
0
8
P2RX7 (AF647)
D
non-MAIT/NKT αβT cells
CD4+
CD8+
Liver
Blood
P2RX7 expression (MFI x10³)
MAIT Vδ2+ Vδ1+ Vδ2-Vδ1- NKT conv. T
CD45RA:
CD27:
CD69:
CD103:
αP2RX7 (AF647)
FMO

**Supplementary Figure S1.**

(A) Boxplots depict *P2RX7* gene transcript levels within the indicated cell types (data source: Gutierrez-Arcelus et al. 2019, boxes show the first to third quartile with median, whiskers encompass 1.5× the interquartile range, and data beyond that threshold indicated as outliers). TPM = transcripts per million. (B) Graph shows mean fluorescence intensity (MFI) of P2RX7 labelling by human blood MAIT cells (MR1-5-OP-RU tetramer$^{+}$CD3$^{+}$), Vδ2$^{+}$ and Vδ2$^{-}$ γδT cells (γδTCR$^{+}$CD3$^{+}$), NKT cells (CD1d-α-GalCer tetramer$^{+}$CD3$^{+}$), and conventional T cells (conv. T; defined as non-MAIT/NKT γδTCR$^{-}$CD3$^{+}$ T cells). A total of 13 donors were analysed across 4 separate experiments. Each symbol represents an individual donor, where upwards- and downwards-pointing triangles represent freshly processed and cryopreserved samples, respectively. Graphs depict individual data points and mean ± SEM. ns P>0.05, *P≤0.05, **P≤0.01 using a Wilcoxon matched pairs signed rank test with a Bonferroni-Dunn correction for multiple comparisons. (C) Histograms depict expression of P2RX7 by CD14$^{+}$FSC$^{hi}$SSC$^{hi}$ cells and CD3$^{+}$ T cells from donors 03 and 04 processed and analysed on the same day. Numbers within histograms represent MFI. FMO = Fluorescence minus one. (D) Graphs depict the MFI of P2RX7 labelling by liver and blood T-cell subsets (blue square symbols) and their respective FMO controls (white circle symbols), as indicated by connecting lines between symbols. A total of 4 human liver donors and 6 blood donors were analysed across 2 separate experiments. The FMO control for NKT cells within one liver donor was not analysed due to low cell numbers. Half-shaded symbols represent matched blood and liver samples from one donor.

### Supp Fig. 2

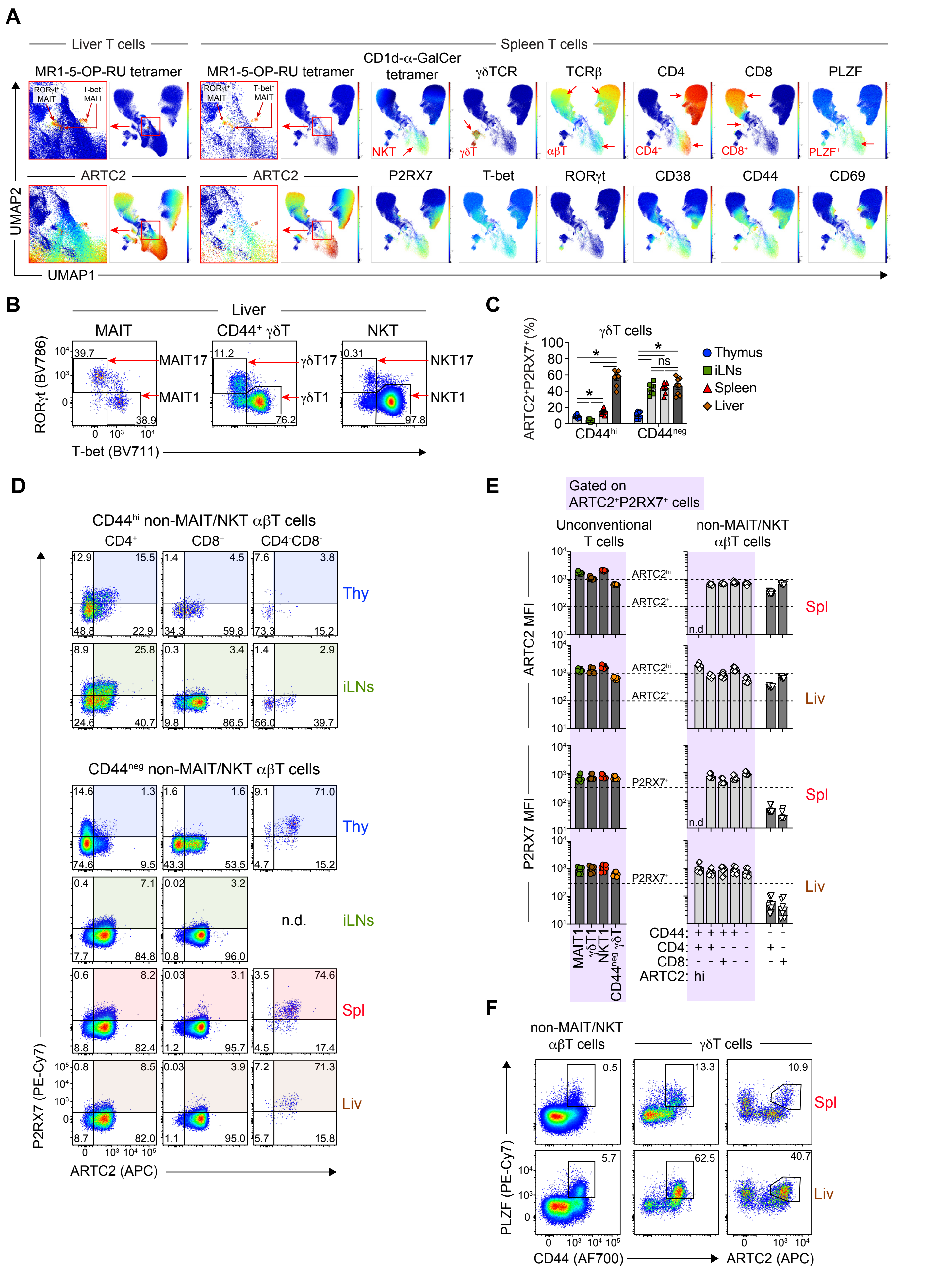

A
Liver T cells
Spleen T cells
MR1-5-OP-RU tetramer
CD1d-α-GalCer tetramer
γδTCR
TCRβ
CD4
CD8
PLZF
ARTC2
P2RX7
T-bet
RORγt
CD38
CD44
CD69
UMAP1
UMAP2
B
Liver
MAIT
CD44+ γδT
NKT
MAIT17
MAIT1
γδT17
γδT1
NKT17
NKT1
RORγt (BV786)
T-bet (BV711)
C
γδT cells
ARTC2+P2RX7+ (%)
Thymus
iLNs
Spleen
Liver
CD44hi
CD44neg
D
CD44hi non-MAIT/NKT αβT cells
CD4+
CD8+
CD4-CD8-
Thy
iLNs
CD44neg non-MAIT/NKT αβT cells
n.d.
Spl
Liv
P2RX7 (PE-Cy7)
ARTC2 (APC)
E
Gated on ARTC2+P2RX7+ cells
Unconventional T cells
non-MAIT/NKT αβT cells
ARTC2 MFI
P2RX7 MFI
CD44:
CD4:
CD8:
ARTC2:
F
non-MAIT/NKT αβT cells
γδT cells
PLZF (PE-Cy7)
CD44 (AF700)
ARTC2 (APC)

**Supplementary Figure S2.**

(A) UMAP representation of flow cytometric analysis of liver and spleen T cells. UMAP plots were generated by concatenation of data from n=2 mice from one of two similar experiments. Red arrows within UMAP plots indicate various T-cell populations. (B) Representative gating of T-bet$^{+}$ROR$\gamma$t$^{-}$ MAIT1, $\gamma\delta$T1, and NKT1 cells, and ROR$\gamma$t$^{+}$T-bet$^{-}$ MAIT17, $\gamma\delta$T17, and NKT17 cells in C57BL/6 WT mouse liver, as indicated by red arrows (C) Graph depicts the percentage of ARTC2$^{+}$P2RX7$^{+}$ cells of CD44$^{neg}$ and CD44$^{hi}$ $\gamma\delta$T cells within C57BL/6 WT mouse organs. n = 3 separate experiments with a total of 8 mice. (D) Flow cytometric analysis of ARTC2 and P2RX7 expression on CD44$^{hi}$ or CD44$^{neg}$ non-MAIT/NKT CD4$^{+}$, CD8$^{+}$, and CD4$^{-}$CD8$^{-}$ $\alpha\beta$T cells. Numbers in FACS plots represent percentage of gated cells. CD44$^{neg}$ CD4$^{-}$CD8$^{-}$ $\alpha\beta$T cells in the iLNs were not determined (n.d.) due to their paucity. (E) Graphs depict the mean fluorescence intensities (MFI) of ARTC2 and P2RX7 expression by the indicated cell types. (F) Flow cytometric analysis of PLZF, CD44, and ARTC2 expression as indicated by non-MAIT/NKT $\alpha\beta$T and $\gamma\delta$T cells from the spleen and liver. (C & E) Graphs depict individual data points and mean ± SEM. With the exception of (F), all mice were injected with the anti-ARTC2 nanobody ‘s+16’ prior to organ harvest.

### Supp Fig. 3

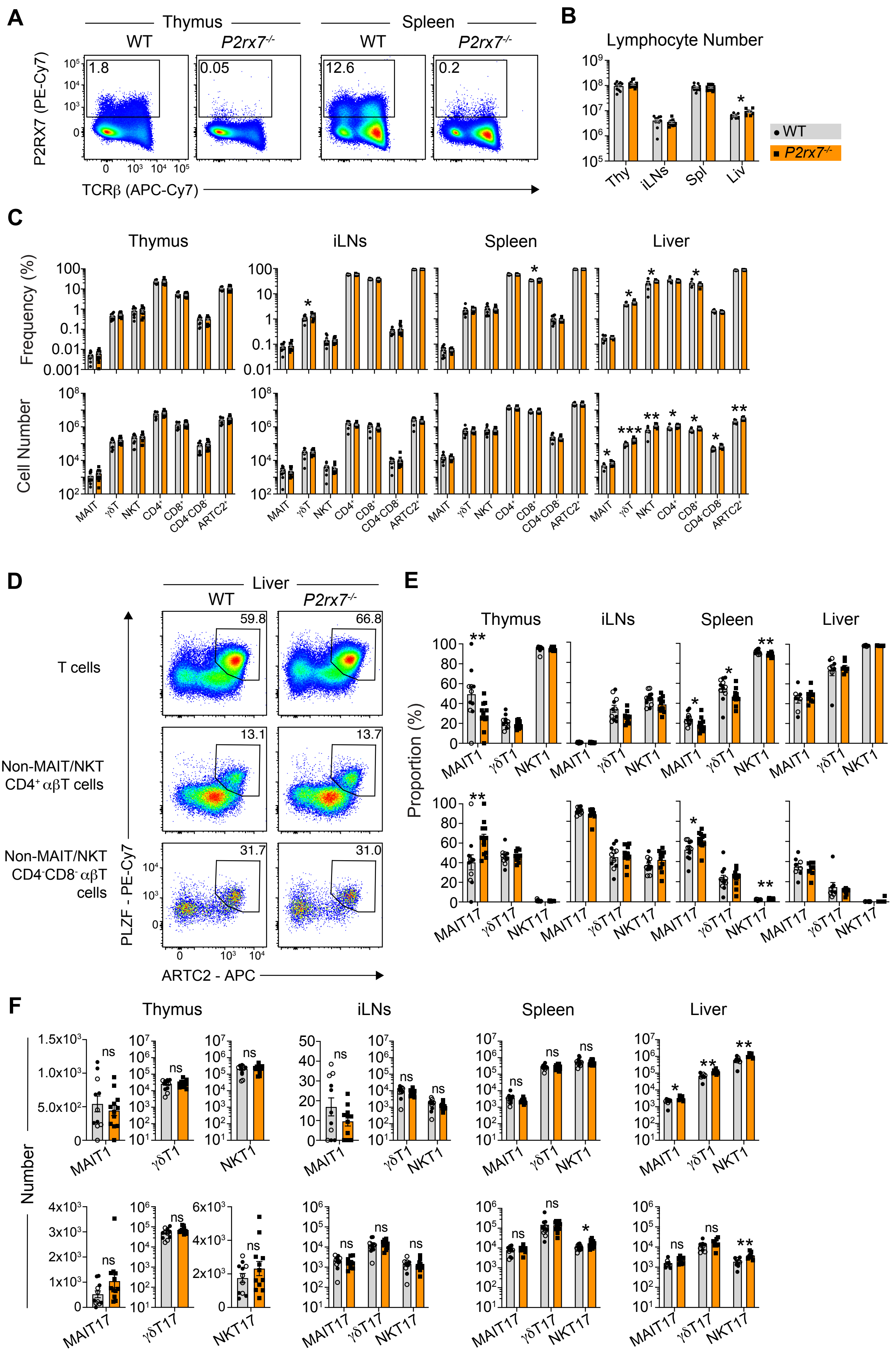
A
Thymus
Spleen
WT
P2rx7-/-
P2RX7 (PE-Cy7)
TCRβ (APC-Cy7)
1.8
0.05
12.6
0.2
B
Lymphocyte Number
Thy
iLNs
Spl
Liv
WT
P2rx7-/-
C
Thymus
iLNs
Spleen
Liver
Frequency (%)
Cell Number
MAIT
γδT
NKT
CD4+
CD8+
CD4-CD8-
ARTC2+
D
Liver
WT
P2rx7-/-
T cells
Non-MAIT/NKT CD4+ αβT cells
Non-MAIT/NKT CD4-CD8- αβT cells
PLZF - PE-Cy7
ARTC2 - APC
59.8
66.8
13.1
13.7
31.7
31.0
E
Thymus
iLNs
Spleen
Liver
Proportion (%)
MAIT1
γδT1
NKT1
MAIT17
γδT17
NKT17
F
Thymus
iLNs
Spleen
Liver
Number
MAIT1
γδT1
NKT1
MAIT17
γδT17
NKT17
ns

**Supplementary Figure S3.**

(A) Flow cytometric analysis of P2RX7 expression on $B220^-$ lymphocytes from the thymus and spleen of WT and *P2rx7*$^{-/-}$ mice. Numbers in FACS plots represent percentage of gated cells. (B) Graph depicts the absolute number of lymphocytes within C57BL/6 WT and *P2rx7*$^{-/-}$ mouse organs. (C) Graphs depict the number and percentage of indicated T-cell populations of total T cells within WT and *P2rx7*$^{-/-}$ mouse organs. $CD4^+$, $CD8^+$, and $CD4^-CD8^-$ T cells are non-MAIT/NKT αβT cells, $ARTC2^+$ T cells are all T cells that express ARTC2. (D) Flow cytometric analysis of PLZF and ARTC2 expression on indicated T-cell populations from the livers of WT and *P2rx7*$^{-/-}$ mice. FACS plots are representative of 8-12 mice per group analysed across n = 2-3 separate experiments. (E & F) Graphs depict the percentage (E) and number (F) of indicated MAIT, γδT, and NKT-cell subsets from specified organs. Open symbols depict WT mice from a different animal house facility used to ensure sufficient numbers for comparison to *P2rx7*$^{-/-}$ mice. (B, C, E, F) Each symbol represents an individual mouse. Graphs depict individual data points and mean ± SEM. n = 2 - 3 separate experiments with a total of 8-12 mice/group. ns P>0.05, *P≤0.05, **P≤0.01, ***P≤0.001 using a Mann-Whitney U test.

### Supp Fig. 4

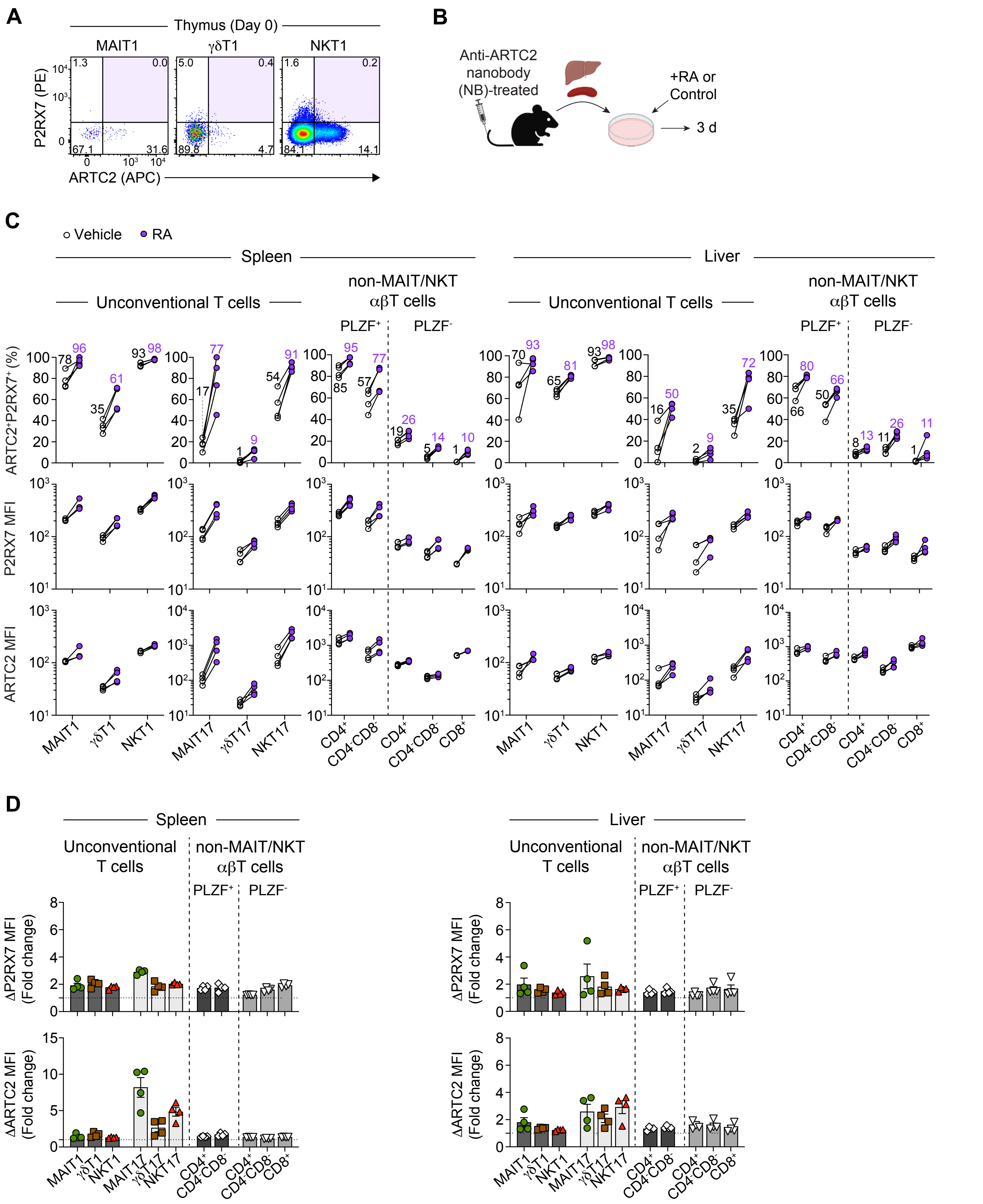

A
Thymus (Day 0)
MAIT1
γδT1
NKT1
P2RX7 (PE)
ARTC2 (APC)
B
Anti-ARTC2 nanobody (NB)-treated
+RA or Control
3 d
C
Vehicle
RA
Spleen
Liver
Unconventional T cells
non-MAIT/NKT αβT cells
PLZF+
PLZF-
ARTC2+P2RX7+ (%)
P2RX7 MFI
ARTC2 MFI
MAIT1
γδT1
NKT1
MAIT17
γδT17
NKT17
CD4+
CD4-CD8-
CD8+
D
Spleen
Liver
Unconventional T cells
non-MAIT/NKT αβT cells
PLZF+
PLZF-
ΔP2RX7 MFI (Fold change)
ΔARTC2 MFI (Fold change)

**Supplementary Figure S4:**

(A) Flow cytometric analysis of ARTC2 and P2RX7 expression by indicated T-cell types from thymus after complement-mediated depletion of immature $CD24^{+}$ thymocytes on (day 0). Numbers within FACS plots represent percentage of gated cells. (B) Experimental schematic. Mouse spleen and livers were cultured for three days in the presence of 20nM all-*trans* retinoic acid (RA) or the vehicle control. Mice were injected with the anti-ARTC2 nanobody (clone: s+16) prior to organ harvest. (C) Graphs show the percentages of $ARTC2^{+}P2RX7^{+}$ cells out of indicated T-cell types from spleen and liver and mean fluorescence intensity (MFI) of P2RX7 and ARTC2 expression. White- and purple-shaded circles represent data from vehicle control or RA-treated cells, respectively. Each symbol represents an individual mouse, where 4 mice were analysed across 2 separate experiments. Connecting lines represent paired data. Numbers above data points represent the average percentage for indicated data sets. (D) Graphs show fold change in P2RX7 and ARTC2 MFI amongst RA-treated cells relative to vehicle controls. Horizontal dotted lines represent a fold change of 1. Graphs depict individual data points and mean ± SEM. n = 2 separate experiments where a total of 4 mice were analysed.

### Supp Fig. 5

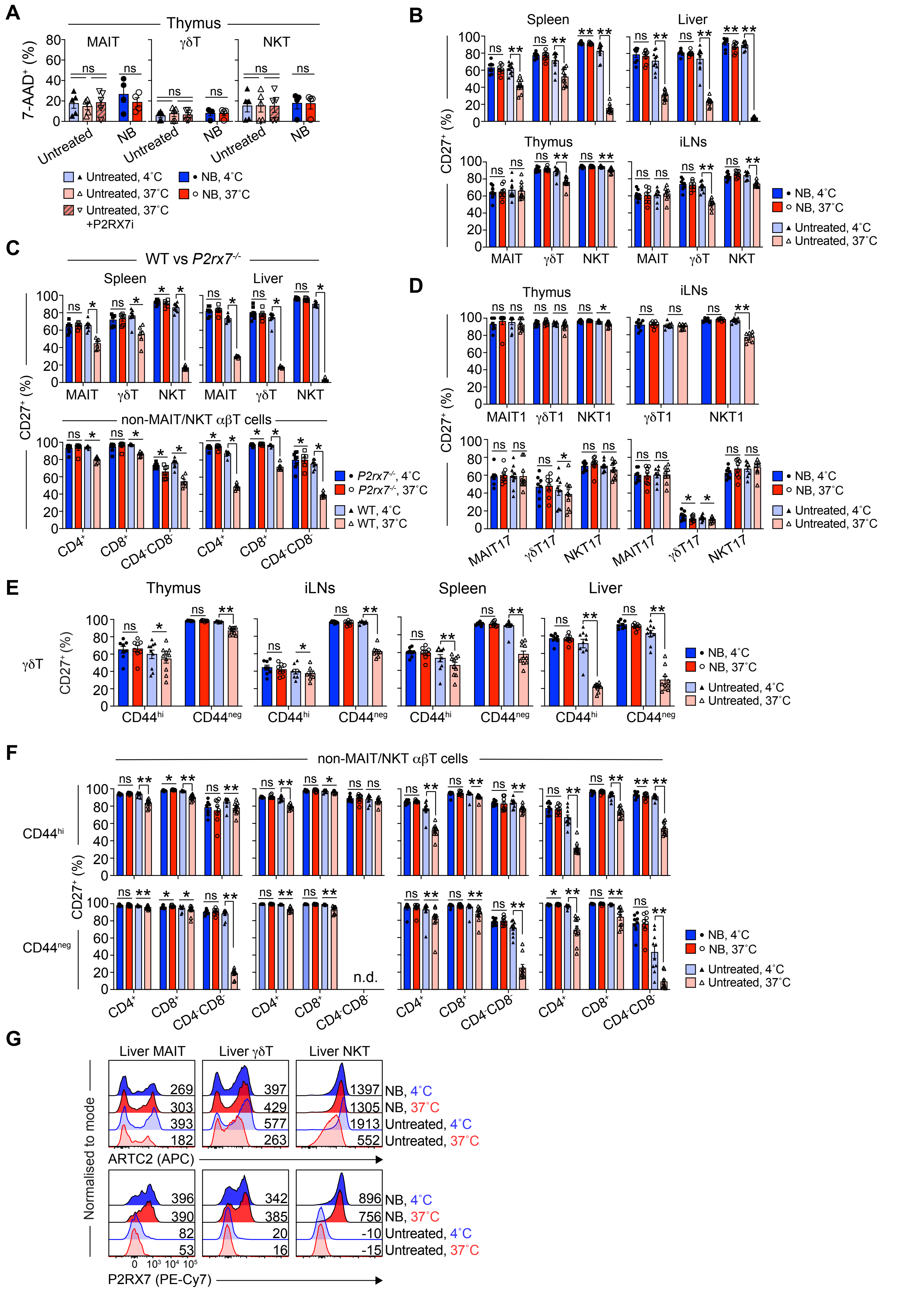
A
Thymus
MAIT
γδT
NKT
7-AAD+ (%)
Untreated
NB
Untreated, 4˚C
Untreated, 37˚C
Untreated, 37˚C +P2RX7i
NB, 4˚C
NB, 37˚C
B
Spleen
Liver
Thymus
iLNs
CD27+ (%)
MAIT
γδT
NKT
C
WT vs P2rx7-/-
Spleen
Liver
non-MAIT/NKT αβT cells
CD4+
CD8+
CD4-CD8-
P2rx7-/-, 4˚C
P2rx7-/-, 37˚C
WT, 4˚C
WT, 37˚C
D
Thymus
iLNs
MAIT1
γδT1
NKT1
MAIT17
γδT17
NKT17
E
Thymus
iLNs
Spleen
Liver
γδT
CD44hi
CD44neg
F
non-MAIT/NKT αβT cells
CD44hi
CD44neg
n.d.
G
Liver MAIT
Liver γδT
Liver NKT
Normalised to mode
ARTC2 (APC)
269
303
393
182
397
429
577
263
1397
1305
1913
552
NB, 4˚C
NB, 37˚C
Untreated, 4˚C
Untreated, 37˚C
396
390
82
53
342
385
20
16
896
756
-10
-15
P2RX7 (PE-Cy7)

**Supplementary Figure S5.**

(A) Graphs depict the percentage of 7-AAD$^+$ MAIT, γδT, and NKT cells from thymus. n = 3 separate experiments with a total of 4-8 mice/group. ns P>0.05, *P≤0.05 using a Mann-Whitney U test, with a correction for multiple comparisons for the untreated groups. Mice were injected with the anti-ARTC2 nanobody (clone: s +16) (NB) or were untreated prior organ harvest. (B – F) Graphs depict the percentage of CD27$^+$ cells out of each indicated cell type from C57BL/6 WT (B, D, E, F), or C57BL/6 WT and *P2rx7*$^{-/-}$ mice (C) after incubation for 30 minutes at 4˚C or 37˚C. Graphs depict individual data points and mean ± SEM. (C) n = 2 separate experiments with a total of 6-8 mice/group. ns P>0.05, *P≤0.05 using a Wilcoxon matched-pairs signed-rank test. (B, D, E, F) n = 3-4 separate experiments with a total of 8-10 mice/group. Each symbol represents an individual mouse. Graphs depict individual data points and mean ± SEM. ns P>0.05, *P≤0.05, **P≤0.01 using a Wilcoxon matched-pairs signed-rank test. (G) Representative overlay histograms of ARTC2 and P2RX7 expression by liver MAIT, γδT, and NKT cells. Dark and light-shaded histograms represent cells from NB-treated and untreated mice, respectively. Numbers within FACS plots represent the mean fluorescence intensity. FACS plots are representative of n = 3-4 separate experiments with a total of 8-10 mice/group. (F) CD44$^{neg}$ CD4$^-$CD8$^-$ αβT cells were not determined (n.d.) in iLNs due to paucity.

Supp Fig. 6

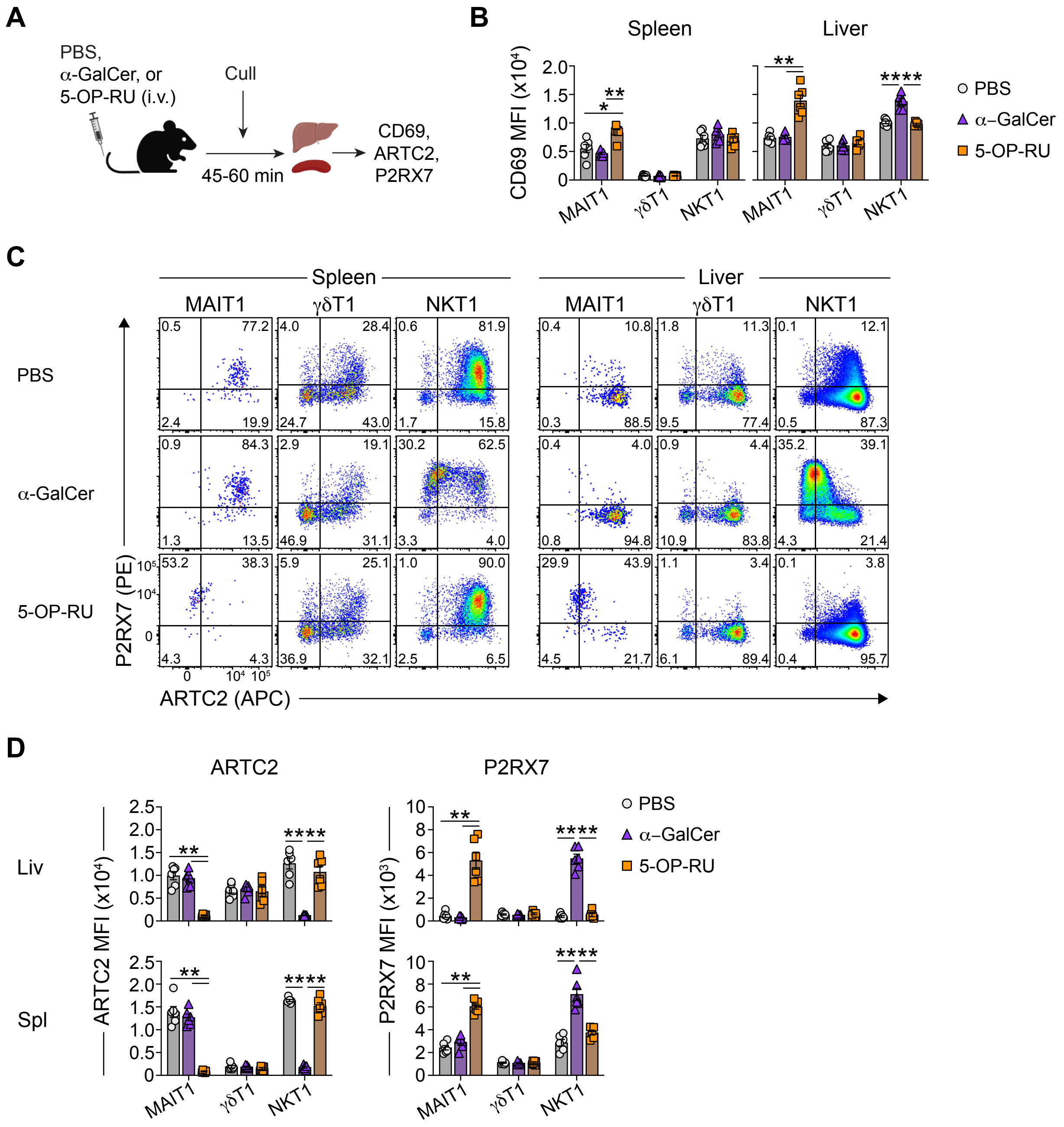
A
PBS,
α-GalCer, or
5-OP-RU (i.v.)
Cull
45-60 min
CD69,
ARTC2,
P2RX7
B
Spleen
Liver
CD69 MFI (x10⁴)
MAIT1
γδT1
NKT1
PBS
α–GalCer
5-OP-RU
C
Spleen
Liver
MAIT1
γδT1
NKT1
PBS
α-GalCer
5-OP-RU
P2RX7 (PE)
ARTC2 (APC)
D
ARTC2
P2RX7
Liv
Spl
ARTC2 MFI (x10⁴)
P2RX7 MFI (x10³)

**Supplementary Figure S6.**

(A) Experimental schematic. Mice were i.v. administered 5-OP-RU (200 pmol), α-GalCer (2μg), or PBS and liver and spleens were harvested 45-60 minutes later. (B & D) Graphs depict the mean fluorescence intensity (MFI) of CD69 (B), ARTC2, and P2RX7 (D) expression by liver and spleen T-bet$^+$ MAIT1, γδT1, and NKT1 cells from indicated treatment groups. ns $P>0.05$ (not shown on graph), *$P\leq0.05$, **$P\leq0.01$ using a Mann-Whitney U test with a Bonferroni-Dunn correction for multiple comparisons. Each symbol represents an individual mouse, where a total of 6 mice per group were analysed across 2 separate experiments. (C) Representative flow cytometric analysis of ARTC2 and P2RX7 expression by the cell types specified. Numbers in FACS plots represent percentage of gated cells.

Supp Fig. 7

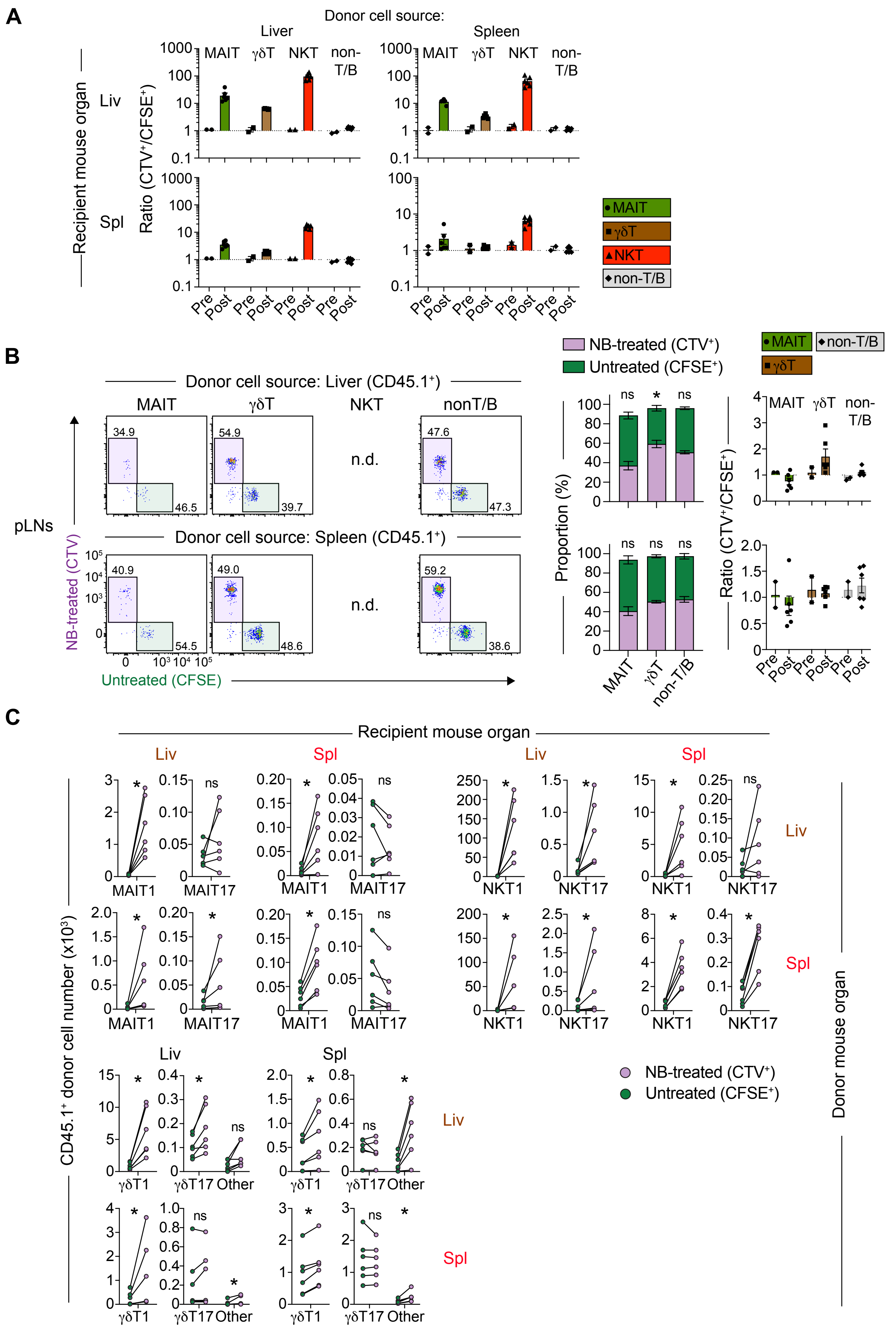
A
Donor cell source:
Liver
Spleen
MAIT
γδT
NKT
non-T/B
Recipient mouse organ
Liv
Spl
Ratio (CTV⁺/CFSE⁺)
Pre
Post
B
NB-treated (CTV⁺)
Untreated (CFSE⁺)
Donor cell source: Liver (CD45.1⁺)
nonT/B
n.d.
pLNs
Donor cell source: Spleen (CD45.1⁺)
NB-treated (CTV)
Untreated (CFSE)
Proportion (%)
ns
*
C
Recipient mouse organ
CD45.1⁺ donor cell number (x10³)
MAIT1
MAIT17
NKT1
NKT17
γδT1
γδT17
Other
Donor mouse organ

**Supplementary Figure S7.**

(A) Graphs show the ratio between percentages of $CTV^+$ cells relative to $CFSE^+$ cells pre- and post-adoptive transfer. (B) Flow cytometric analysis of donor $CD45.1^+$ MAIT, γδT, and non-T/B cells sourced from liver or spleen and recovered from the pooled peripheral lymph nodes (pLNs) of recipient mice 8 days after adoptive transfer. MAIT and γδT-cell FACS plots were generated by concatenation of data from all (n = 3) mice of the same group from one of two similar experiments. NKT cells were not analysed due to low lymph node cell numbers. Stacked bar charts depict the percentage of recovered donor $CTV^+$ and $CFSE^+$ cells sourced from liver and spleen after adoptive transfer. ns P>0.05, *P≤0.05, **P≤0.01 using a Wilcoxon matched-pairs signed-rank test for NB-treated vs untreated. Graph shows ratio between percentage of $CTV^+$ cells relative to $CFSE^+$ cells pre- and post-adoptive transfer. (A & B) Graphs depict individual data points and mean ± SEM. Post: each symbol represents an individual mouse, where a total of 6 mice/group were analysed across n = 2 separate experiments. Pre: each symbol represents a separate experiment where cells from 8 mice were pooled. (C) Graphs depict the absolute numbers of recovered $CTV^+$ and $CFSE^+$ MAIT, NKT, and γδT-cell subsets after adoptive transfer. MAIT1, γδT1, and NKT1 cells defined as $CD44^+CD319^+$. MAIT17 and NKT17 cells defined as $ICOS^+CD319^-$. γδT17 cells defined as $CD44^{hi}CD319^-$. Remaining $CD44^-CD319^-$ γδT cells defined as 'other'. Connecting lines represent paired data (cells recovered from the same organs). ns P>0.05, * P≤ 0.05 using a Wilcoxon matched-pairs signed-rank test.

Supp Fig. 8

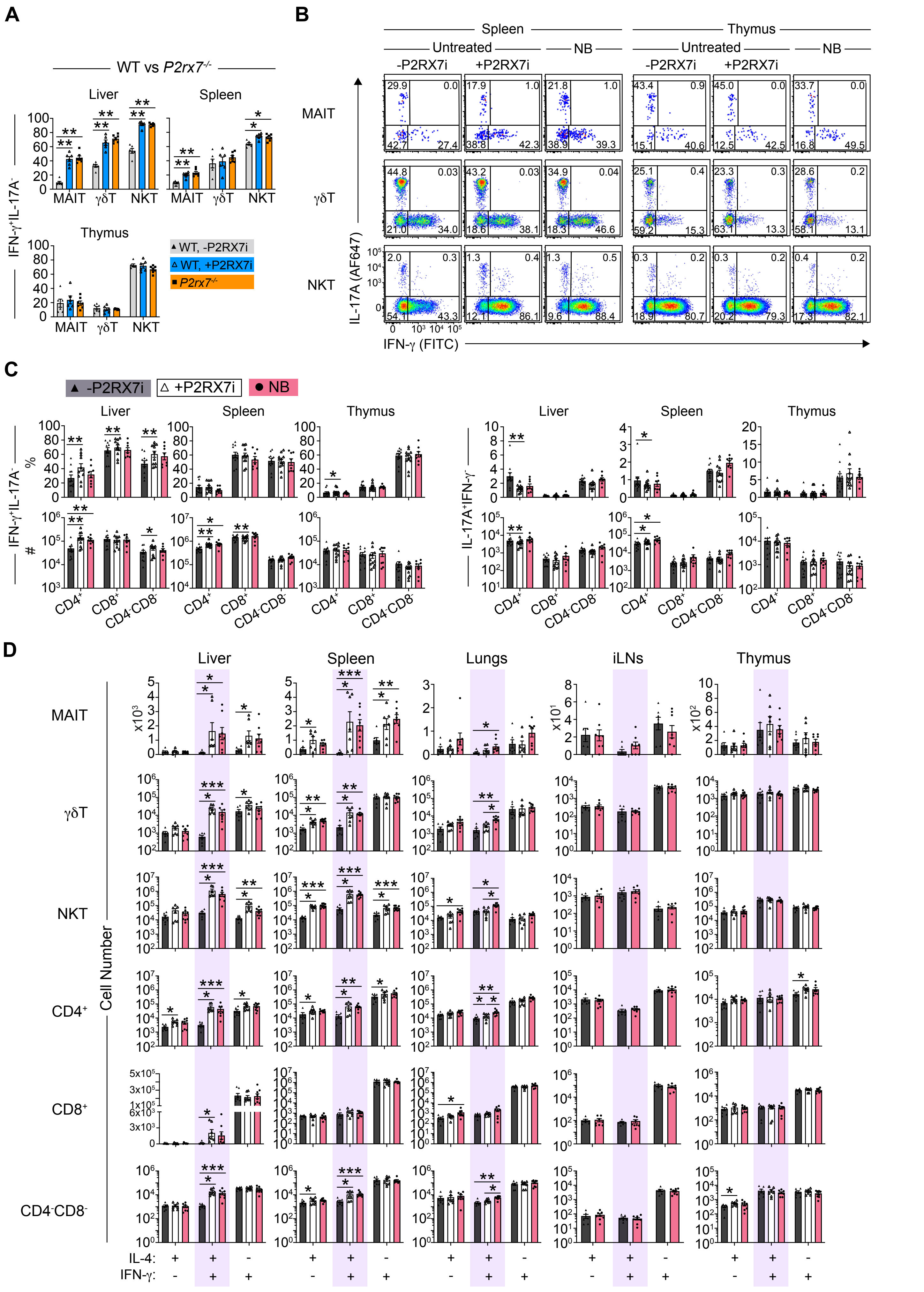
A
WT vs P2rx7-/-
Liver
Spleen
Thymus
IFN-γ+IL-17A-
MAIT
γδT
NKT
WT, -P2RX7i
WT, +P2RX7i
P2rx7-/-
B
Spleen
Thymus
Untreated
NB
-P2RX7i
+P2RX7i
IL-17A (AF647)
IFN-γ (FITC)
C
▲ -P2RX7i
△ +P2RX7i
● NB
%
#
IL-17A+IFN-γ-
CD4+
CD8+
CD4-CD8-
D
Liver
Spleen
Lungs
iLNs
Thymus
Cell Number
IL-4:
IFN-γ:

**Supplementary Figure S8.**

(A) Graphs depict percentages of IFN-$\gamma^{+}$IL-17$^{-}$ cells from C57BL/6 WT and *P2rx7*$^{-/-}$ mice after stimulation for 4 hours in the presence of PMA and ionomycin, with or without the P2RX7 inhibitor (P2RX7i), A438079 (10μM). n.s. P>0.05 (not shown on graph), *P≤0.05, and **P≤0.01 using a Mann-Whitney U test with a Bonferroni-Dunn correction for multiple comparisons. (B) Flow cytometric analysis of IL-17A and IFN-γ expression by CD44$^{+}$ MAIT, γδT, NKT cells from spleen and thymus after PMA/ionomycin stimulation. Numbers in FACS plots represent percentage of gated cells. Mice were treated the anti-ARTC2 nanobody (clone: s+16) (NB) or were left untreated prior to organ harvest. Untreated mouse cells were stimulated with or without the P2RX7i. (C & D) The percentage (%) and absolute number (#) of IFN-$\gamma^{+}$IL-17A$^{-}$ and IL-17A$^{+}$IFN-$\gamma^{-}$ non-MAIT/NKT αβT-cell subsets (C), and the absolute number for each IL-4/IFN-γ subset of CD44$^{+}$ MAIT, γδT, NKT cells, and of non-MAIT/NKT αβT-cell types (D) were graphed. (C) n = 3-5 separate experiments with a total of 8-12 mice/group. (D) n = 3 separate experiments with a total of 7-8 mice/group. n.s. P>0.05 (not shown on graph), *P≤0.05, **P≤0.01, ***P≤0.001, and ****P≤0.0001 using a Wilcoxon matched-pairs signed-rank test for +P2RX7i vs -P2RX7i or a Mann-Whitney U test with a Bonferroni-Dunn correction for multiple comparisons for all other comparisons between conditions.

### Supp Fig. 9

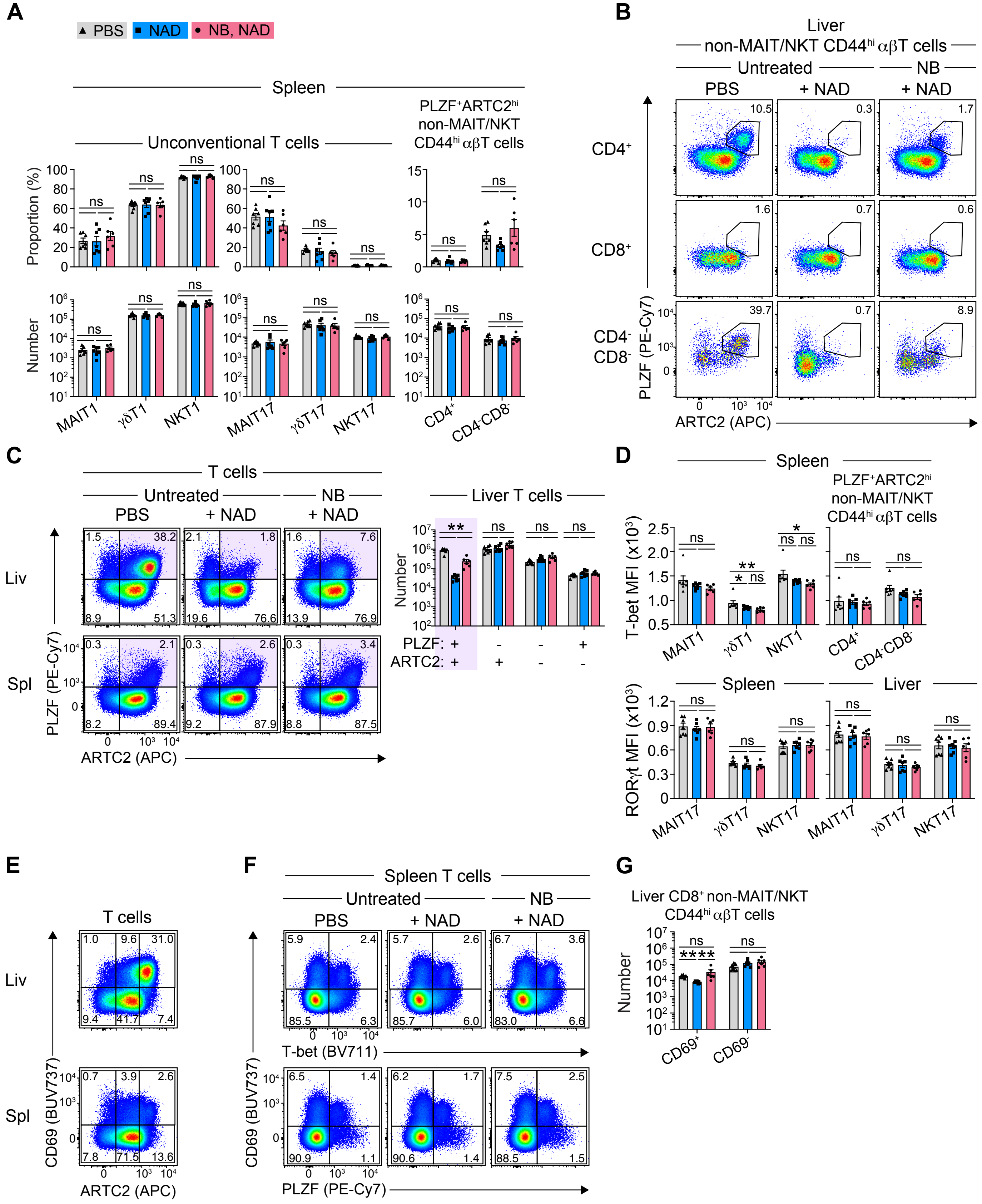

A
PBS
NAD
NB, NAD
Spleen
Unconventional T cells
PLZF+ARTC2hi non-MAIT/NKT CD44hi αβT cells
Proportion (%)
Number
ns
MAIT1
γδT1
NKT1
MAIT17
γδT17
NKT17
CD4+
CD4-CD8-
B
Liver
non-MAIT/NKT CD44hi αβT cells
Untreated
NB
PBS
+ NAD
CD4+
CD8+
CD4- CD8-
PLZF (PE-Cy7)
ARTC2 (APC)
10.5
0.3
1.7
1.6
0.7
0.6
39.7
0.7
8.9
C
T cells
Liv
Spl
Liver T cells
PLZF:
ARTC2:
**
D
T-bet MFI (x10³)
RORγt MFI (x10³)
*
E
CD69 (BUV737)
F
Spleen T cells
T-bet (BV711)
G
Liver CD8+ non-MAIT/NKT CD44hi αβT cells
****
CD69+
CD69-

**Supplementary Figure S9.**

(A) Anti-ARTC2 nanobody (clone: s+16) (NB)-treated or untreated mice were intravenously injected with PBS or nicotinamide adenine dinucleotide (NAD) 30 minutes prior to organ harvest. Graphs depict the percentage and numbers of the indicated cell types from spleen and livers of specified mice. (B & C) Flow cytometric analysis of PLZF and ARTC2 expression by non-MAIT/NKT αβT-cell subsets (B) and by all T cells from liver and spleen (C). Graph in (C) depicts the absolute numbers of the indicated PLZF/ARTC2 subsets from liver. (D) Graphs depict the mean fluorescence intensity (MFI) of T-bet and RORγt expression. (E & F) Flow cytometric analysis of CD69 and ARTC2 (E), T-bet, and PLZF (F), as indicated, by liver and spleen T cells. (G) Graph depicts the absolute numbers of $CD69^+$ and $CD69^-$ $CD44^{hi}CD8^+$ non-MAIT/NKT αβT cells from livers of indicated mice. (A, C, D, G) Graphs depict individual data points and mean ± SEM. n = 2 separate experiments with a total of 6-7 mice/group. Each symbol represents an individual mouse. n.s. P>0.05, *P≤0.05, **P≤0.01 using a Mann-Whitney U test with a Bonferroni-Dunn correction for multiple comparisons. (B, C, E, F) Numbers in FACS plots represent percentage of gated cells.
